# A *multiple-causal-gene-cluster* model underlying GWAS signals of Alzheimer’s disease

**DOI:** 10.1101/2021.05.14.444131

**Authors:** Min Xu, Qianjin Liu, Rui Bi, Yu Li, Chunhua Zeng, Zhongjiang Yan, Quanzhen Zheng, Xiao Li, Chunli Sun, Maosen Ye, Xiong-Jian Luo, Ming Li, Deng-Feng Zhang, Yong-Gang Yao

**Affiliations:** Key Laboratory of Animal Models and Human Disease Mechanisms of the Chinese Academy of Sciences & Yunnan Province, and KIZ/CUHK Joint Laboratory of Bioresources and Molecular Research in Common Diseases, Kunming Institute of Zoology, Chinese Academy of Sciences, Kunming 650204, China; Kunming College of Life Science, University of Chinese Academy of Sciences, Kunming 650204, China; Center for Excellence in Animal Evolution and Genetics, Chinese Academy of Sciences, Kunming 650204, China; CAS Center for Excellence in Brain Science and Intelligence Technology, Chinese Academy of Sciences, Shanghai 200031, China

**Keywords:** Alzheimer’s disease, GWAS, functional genomics, multi-omics, fine-mapping, functional variant, causal gene

## Abstract

Genome-wide association studies (GWASs) have identified dozens of genetic susceptibility loci for Alzheimer’s disease (AD). Nevertheless, the underlying causal variants and biological mechanisms remain elusive. Here, we systematically integrated AD GWAS with comprehensive multi-omics data, and distilled 304 potentially functional variants and 166 causal genes from 49 loci. Intriguingly, we found that most of AD GWAS loci contain multiple functional variants and causal genes. *In vitro* assays showed that one functional variant regulated multiple genes in the 11p11.2 locus (the CELF1/SPI1 locus) and alteration of these target genes contributed to AD-related molecular processes, supporting the co-existence of multiple functional variants and AD-relevant causal genes within a single locus. We thus proposed a *multiple-causal-gene-cluster* model that co-dysregulation of a cluster of genes within a single GWAS loci individually or synergistically contribute to AD development. This model provides a novel insight into the biological mechanisms underlying the GWAS loci of complex traits.

## Introduction

Alzheimer’s disease (AD) is the most common cause of dementia with a genetic heritability of 58–79% ^1–3^. Genome-wide association studies (GWASs) have identified dozens of genetic loci (tagged by single nucleotide polymorphisms [SNPs]) contributing to susceptibility of sporadic AD ^4–9^. The recognition of such risk loci has provided a rich resource for furthering our understanding of the pathogenesis of AD. However, it is challenging to elucidate the genetic mechanisms underlying these tagged-SNP based GWAS signals ^4, 10–16^. The large number of non-coding variants with unknown function, the complex linkage disequilibrium (LD) among variants, and the complicated regulatory structures all impede the inference of disease-causing variants and their biologically-relevant effect genes, and the depicting of complete pathogenic mechanisms ^4, 10^. In current practice, a variant affecting regulatory activity (usually represented by expression Quantitative Trait Loci [eQTL]) is usually assigned as the *causal variant* within the corresponding GWAS loci, and the gene (or genes) whose expression is associated or regulated by the variant is assigned as the *causal gene* ^5, 6, 17^. However, this kind of brief inference is greatly influenced by LD among SNPs and expressional correlations among genes. Systematic investigations with multi-dimensional types of omics data ^9, 18–20^, together with experimental assays are essential to improve the decipherment of AD GWAS signals . In addition to the fine-mapping of functional variants/genes, a more challenging issue might be to look at the mechanistic model by which the functional variants and their effect genes underlying GWAS signals contribute to AD.

In this study, we performed a systematic functional genomic study of the causal/functional variants and their biologically relevant effect genes, as well as the regulatory mechanisms within the AD GWAS risk loci. Summary statistics of AD GWAS were integrated with large-scale multi-omics data including histone modification, open chromatin, transcription factor (TF) binding, eQTL, single-cell differential expression profiling, and three-dimensional (3D) chromatin interaction data from AD relevant tissues or cells. The fine-grained and unbiased annotation distilled 304 potentially functional variants that might disrupt regulatory activities from 7,884 AD-associated SNPs within 49 GWAS risk loci. A total of 166 genes might be regulated by these functional variants and had expressional changes at the early stage of AD, which were supposed to be causal genes of AD. Intriguingly, we observed a pattern of multiple functional variants and causal genes co-existing within a single GWAS locus for many loci, especially for the 11p11.2 locus (the CELF1/SPI1 locus). Functional variants in the 11p11.2 could be validated by cellular assays and these variants synergistically regulated a cluster of causal genes via a variety of regulation models including chromatin interactions. Co-dysregulation of causal genes in the 11p11.2 locus influenced several AD-related pathways, and finally contributed to the molecular processes relevant to AD. This “one GWAS signal, multiple-functional-variants, multiple-causal-gene-cluster” pattern was not restricted to AD, but was also shared by GWAS loci of other complex traits and diseases. Our study provided a paradigm for elucidating the complex biological mechanism of complex traits in the post-GWAS era.

## Results

### Multi-omics-based systematic functional genomic study strategy for the inference of functional variants and causal genes within AD GWAS loci

The AD-associated common variants (minor allele frequency [MAF]>0.01) with genome-wide significance (GWAS *P*<5x10^-8^) or sub-threshold significance (GWAS *P*<1x10^-5^) and SNPs in linkage with these variants (R^2^>0.8) from three large-scale GWASs (the Lambert study ^8^, the Kunkle study ^5^, and the Jansen study ^6^) were used for inference of functional variants and causal genes. Functional variants were defined as those located in actively regulatory elements (promoters or enhancers) and could affect transcriptional events. Multiple levels of omics data were integrated to prioritize potentially functional variants (**Fig. 1A**). Briefly, histone modification data of AD-related tissues or cells (i.e. brain tissues, neural cells, and monocytes; **Supplementary Table S1** ^21, 22^) were used to identify variants located in promoters (H3K4me3 and H3K9ac) ^23^ or enhancers (H3K4me1 and H3K27ac) ^24^. Open chromatin data of AD-related tissues or cells ^25–27^ were used to determine whether regulatory elements were transcriptionally active. Transcription factor (TF) binding data (**Supplementary Table S2**, ^21, 22^) integrated with the atSNP algorithm ^28^ were used to predict whether regulatory elements containing different alleles of candidate variants had different binding affinities with TFs. The biological consequence of these regulatory variants were measured by integrating the eQTL data of brain ^7, 29^ and monocytes ^30, 31^ which aimed at identifying variants with a potential to affect expression of nearby genes. These functional variants regulated eQTL genes were further tested whether their dysregulations are disease relevant. Since the expressional alteration of a gene at the early stage of the disease is more likely to be caused by upstream events such as regulatory genetic variations ^32^, we therefore defined those genes regulated by functional variants and differentially expressed in single-cell RNA sequencing data of brain tissues from the early stage of AD ^33^ as *causal* genes in this study (**Fig. 1A**).

**Figure 1.**
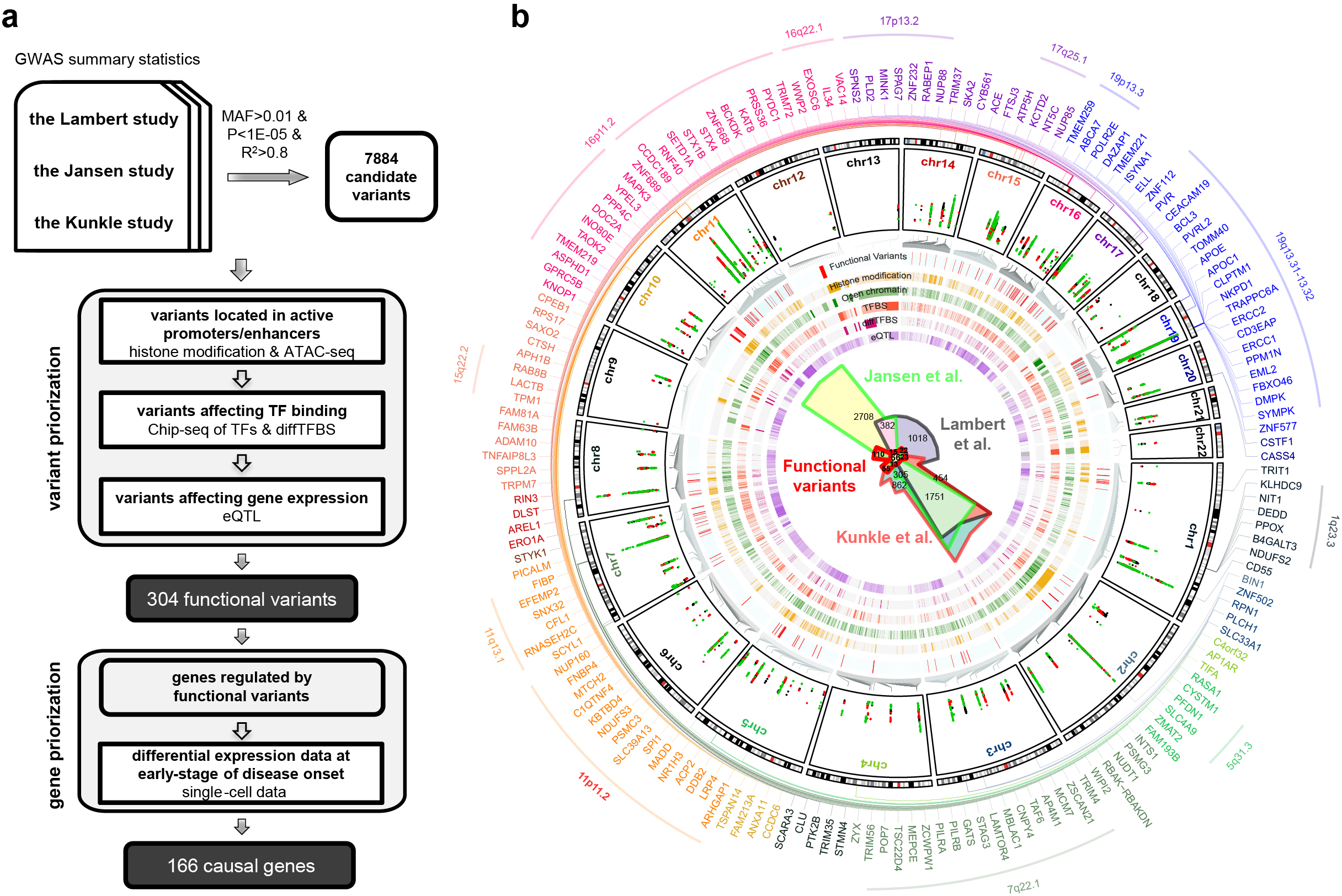
Systematic functional annotation of AD GWAS loci based on multi-omics data. **a. Workflow of functional annotation of AD GWAS loci.**MAF: minor allele frequency; P: GWAS *P*-value; ATAC-seq: assay for transposase-accessible chromatin with high throughput sequencing; TFs: transcriptional factor; diffTFBS: differential TF binding; eQTL: expressional quantitative loci. **b. Genome-wide view of distributions of prioritized functional variants and effect genes.** The height of the plot in GWAS circle indicates the significance of the association (-log10 P-value), colors of dots indicated different GWAS studies: green for the Jansen study, black for the Lambert Study, and red for the Kunkle study. The colored line in each circular band indicated SNP at this site had positive signals in the corresponding annotation. GWAS loci with more than 3 effect genes were highlighted with arcs.

### Functional annotation identified 304 functional variants and 166 causal genes accounting for AD GWAS signals

A total of 7,884 candidate variants within 49 GWAS loci distilled from the three GWASs were subjected to a systematic and unbiased functional annotation. Among these candidate variants, 304 potentially functional variants were recognized (**Fig. 1B**, **Supplementary Table S3**), which regulated 166 potentially AD causal genes that were differentially expressed in brain of early-stage AD (**Fig. 1B**, **Supplementary Table S4**). These causal genes were significantly enriched in several known AD-related pathways (Gene ontology, biological process, FDR<0.05) ^32, 34–39^, such as the production and clearance of amyloid-beta (Aβ), energy metabolism, glia cell activation and neuroinflammation, neuron projection and synaptic transmission, endosome and lysosome-related pathways, lipid metabolism, and infection **(Supplementary Fig. S1)**. Such fine-grained and unbiased resources might benefit the community for further functional investigation, we thus deposited the detailed annotations of functional variants and causal genes in our webserver AlzData (http://www.alzdata.org/functional_SNP_1.php) (**Supplementary Fig. S2**).

### Multiple functional variants and causal genes co-existed in one GWAS locus

Among the 304 functional variants and 166 causal genes within the 49 risk loci (Fig. 1B), we found that multiple functional variants and causal genes, rather than a single causal variant/genewere accounting for the GWAS signals for most of the loci. For example, 1q23.3 (the *ADAMTS4* locus), 5q31.3 (the *HBEGF* locus), 7q22.1 (the *PILRA* locus), 11p11.2 (the *CELF1*/*SPI1* locus), 11q13.1 (the *FIBP* locus), 15q22.2 (the *APH1B* locus), 16p11.2 (the *KAT8* locus and the *FAM57B* locus), 16q22.1 (the *IL34* locus), 17p13.2 (the *SCIMP* locus), 17q25.1 (the *SLC16A5* locus), 19p13.3 (the *ABCA7* locus), and 19q13.31-13.32 (the *APOE* locus) all harbored more than three potentially causal genes (**Fig. 1B**, **Fig. 2A**, **Supplementary Fig. S3-S13**). These loci had very complex LD structures, and SNPs within these loci were distributed in several independent linkage blocks (**Supplementary Fig. S3-S13**, **Fig. 2A**), which suggested that the association signals at each locus could not be the result of a single variant. Loci 7q22.1 (**Supplementary Fig. S5**), 11p11.2 (**Fig. 2A**), 16p11.2 (**Supplementary Fig. S8**), and 19q13.31-13.32 (**Supplementary Fig. S13**) had a dozen or dozens of potentially functional variants and causal genes. For instance, 63 functional variants and 19 causal genes were prioritized for the 19q13.31-13.32 locus (**Supplementary Fig. S13**), and 24 functional variants and 15 causal genes were identified for the 11p11.2 locus (**Fig. 2A**). This co-existence of multiple functional variants and causal genes constituted a gene-cluster pattern, which suggested a synergistic mode of regulation of these genes/functional variants.

**Figure 2.**
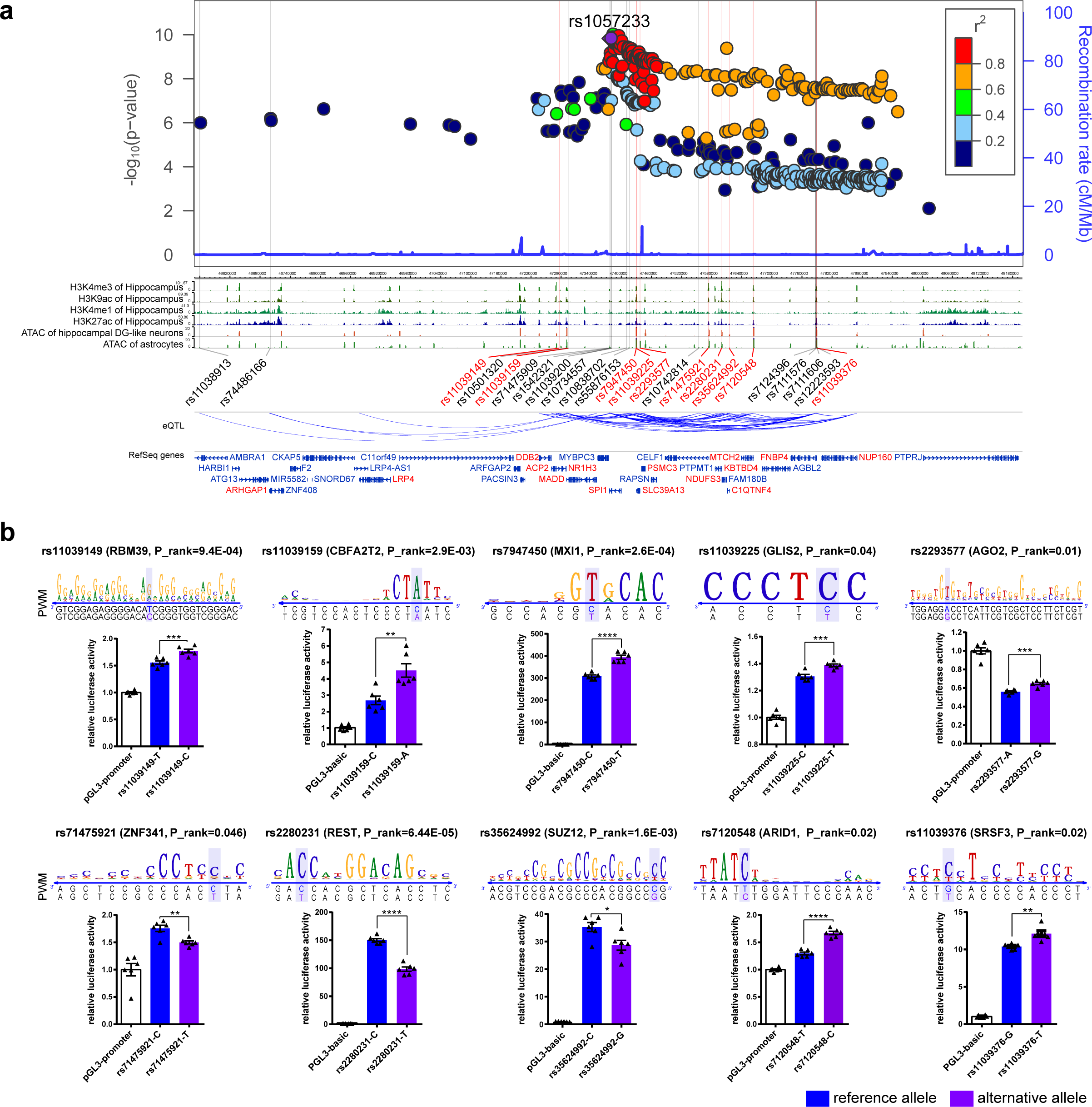
Validation of the existence of multiple functional variants in the 11p11.2 locus. **a. Functional variants and effect genes in the 11p11.2 locus. (***Upper panel*) locus plot of GWAS results from the Kunkle study; (*Lower panel*) functional annotation data including histone modification data from hippocampus, assay for transposase-accessible chromatin with high throughput sequencing (ATAC-seq) data from neurons and astrocytes, and eQTL data from brains and monocytes showing associations among functional variants and effect genes; functional variants were highlighted with grey lines, and effect genes were highlighted with red; variants marked in red were those validated by luciferase assays. **b. Luciferase assays showing functional variants in the 11p11.2 locus in 293T cells.** The binding motif (PWM) of the TF (*upper panel*), and result of luciferase assay of the target variant in 239T cell (*lower panel*) were shown for each variant. Shown results were representative of three independent experiments with similar results. Values are presented as mean ± SD (n = 6 replicates) and are measured by two-sided student’s *t* test. *, *P*<0.05; **, *P*<0.01; ***, *P*<0.001; ****, *P*<0.0001.

### Cellular validation of the functional-variant-cluster in the 11p11.2 locus

We took the 11p11.2 locus (the *CELF1*/*SPI1* locus), which was one of the most reliable and complicated AD risk loci reported so far ^5, 6, 8^, as an example to verify the existence of multiple functional variants/genes. The 11p11.2 locus contained over 450 AD-associated SNPs which extended for almost 1.5 million bases (Mb, chr11: 46.5M-48M), and 24 potentially functional variants and 15 causal genes were predicted for this locus (**Fig. 2A**). Three long linkage blocks (LBs, *R^2^*>0.8) were discerned at this locus, with 170, 74, and 160 SNPs for LB1, LB2, and LB3, respectively (**Supplementary Table S5**). Haplotype constituted by the alternative alleles of the functional variants in LB1 was associated with a low risk for AD (GWAS beta<0), while those of LB2 and LB3 were associated with an increased AD risk (GWAS beta>0). A number of potentially causal genes have been reported for this locus in the previous studies, e.g. *MTCH2*, *SPI1*, *CELF1*, and *PSMC3* ^5, 17, 40^. However, due to the complex linkage structure and eQTL signals, the exact causal variants and genes in this locus have remained unknown ^5, 17, 40–42^. In order to test the reliability of these functional variants in 11p11.2, we performed luciferase reporter assay for 10 out of the 24 functional variants (marked in red in **Fig. 2A**) in HEK293T cells. Consistent with the regulatory element annotations, different alleles of all these variants could affect luciferase reporter activities (**Fig 2B**), which confirmed the co-existence of multiple functional variants in the 11p11.2 locus.

### One functional variant could regulate expressions of multiple potentially causal genes in the 11p11.2 locus

Complicated eQTL associations were observed among the identified functional variants and causal genes in the 11p11.2 locus (**Fig. 3A**). For instance, the AD-protective allele of rs2293577 (G) was associated with higher expression of most of the causal genes in the 11p11.2 region (except for *SPI1*, *C1QTNF4*, and *NUP160*), whereas the risk allele of rs2280231 (A) had an opposite effect on several genes (**Fig. 3A**). To dissect the regulation pattern of the functional-variant-cluster, we tested whether one functional variant could regulate expression of multiple genes or one gene was regulated by several variants (**Fig. 3A**) by cellular assays. We performed base-editing on two functional variants rs2293577 and rs2280231 in HEK293T cells (**Supplementary Fig. S14A-B**), and analyzed the mRNA expression of their potential target genes as revealed by eQTL analysis (**Fig. 3A**). Indeed, quantification of mRNA expression of most of potential target genes of these two functional variants were significantly altered in cells carrying the different alleles (**Fig. 3B**), which supported the notion that one functional variant could regulate multiple causal genes and one causal gene could be regulated by several functional variants. Note that the expression alteration patterns of *ACP2* and *NUP160* were different from that predicted by eQTL, and this might be caused by the tissue heterogeneity.

**Figure 3.**
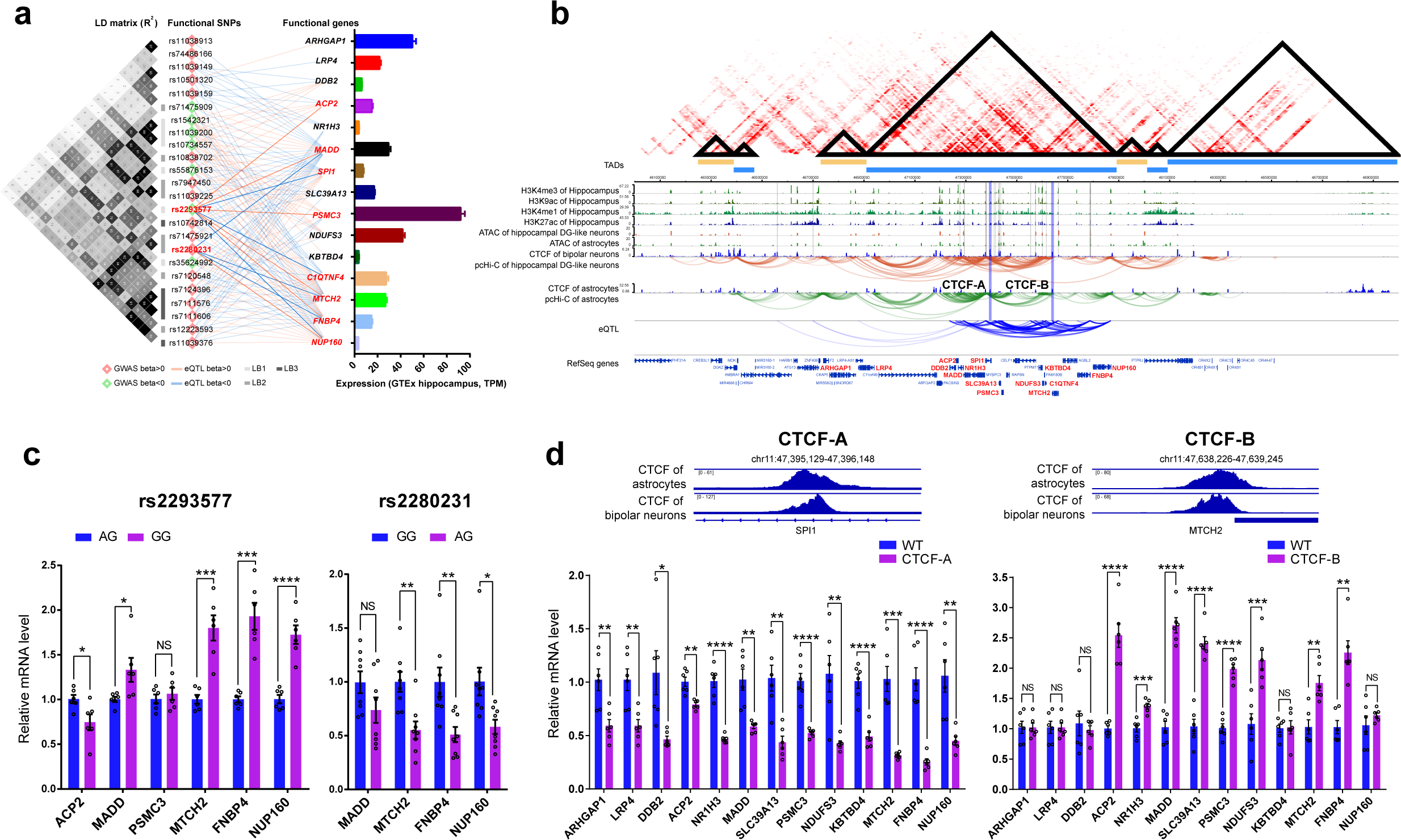
Chromatin interactions mediated the complex regulation of functional variants on effect genes in the 11p11.2 locus. **a. Effect of functional variants in the 11p11.2 locus on expression of multiple functional genes.** (*Left panel*) the linkage pattern among the 24 functional variants represented by R^2^; (*Right panel*) the eQTL relationship among 24 functional variants and 15 functional genes, lines linked functional variants and genes represented eQTL effect, with colors representing the direction of eQTL. The direction of GWAS association was also marked by different colors. **b. Base-editing for rs2293577 and rs2280231 in 293T cells validated the notion that one functional variant could affect expression of multiple functional genes.** Values are presented as mean ± SD (n = 8 cell clones for each genotype of rs2293577; n=6 cell clones for each genotype of rs2280231). **c. Regulatory elements harboring functional variants or those of functional genes interacted with each other via chromatin loops in the 11p11.2 locus.** (*Upper panel*) Chromatin interaction heatmap for the chr11:46M∼49M region, TADs were highlighted with triangles and orange or blue bars; (*lower panel*) functional annotation data including histone modification data from hippocampus, assay for transposase-accessible chromatin with high throughput sequencing (ATAC-seq) data, CTCF chromatin immunoprecipitation followed by sequencing (Chip-seq) data, promoter-caputure high-throughput chromosome conformation capture (pcHiC) data for neurons and astrocytes, and eQTL data from brains and monocytes presenting the association among functional variants and effect genes. Functional variants were highlighted with grey lines, and effect genes were highlighted with red; CTCF-binding sites for CRISPR/Cas9 knockout were highlighted with blue bars. **d. Knockout of two key CTCF-binding sites mediating frequent chromatin interactions in the 11p11.2 locus by CRISPR/Cas9 significantly affected expression of multiple genes. (***Upper panel*) CTCF-binding sites for CRISPR/Cas9 knockout represented by Chip-seq data of CTCF from astrocytes and neurons; (*lower panel*) Quantification of mRNA level of effect genes in wild-type controls (WT) and CTCF-binding site knockout cells (CTCF-A or CTCF-B). *, *P*<0.05; **, *P*<0.01; ***, *P*<0.001; ****, *P*<0.0001; NS, not significant, *P*>0.05; two-sided student’s *t* test. Values are presented as mean ± SD (n = 6 replicates).

### Chromatin interactions mediated the complex regulatory relationships among functional variants and causal genes in the 11p11.2 locus

Chromatin interaction data from hippocampal dentate gyrus (DG)-like neurons showed that the genomic region containing most functional variants and causal genes in the 11p11.2 locus was in one topologically-associated domain (TAD) (**Fig. 3C**). Intriguingly, there were frequent chromatin interactions among regulatory elements harboring functional variants and promoters/enhancers of causal genes, and the pattern was highly consistent with the eQTL association signals (**Fig. 3C**). This observation indicated that a large proportion of eQTL signals in this locus were mediated by chromatin interactions and one functional variant might regulate its proxy or distal target genes via chromatin-loops. In order to validate this speculation, we knocked out the CTCF-binding sites that mediated frequent chromatin interactions in the 11p11.2 locus (marked in **Fig. 3C**) in a U251 glia cell line with a stable expression of mutant APP (APP-K670N/M671L) constructed in our previous study (U251-APP cell) ^43, 44^. Two CTCF-binding sites were successfully deleted in two cell clones by CRISPR/Cas9, one was in the intron of *SPI1* (CTCF-A cell) and another one in the intergenic region near the 3’UTR of *MTCH2* (CTCF-B cell) (**Fig. 3D** and **Supplementary Fig. S14C-D**). Potential off-target sites were checked by Sanger sequencing and only one intronic off-target site was identified (**Supplementary Fig. S14E-N**). mRNA expression levels of most of causal genes in the 11p11.2 locus were concurrently affected by the CTCF-binding site knockout, with 13 genes downregulated in CTCF-A cells, and eight genes upregulated in CTCF-B cells **(Fig. 3D)**. This result proved the complex regulation pattern among functional variants and causal genes in the 11p11.2 locus was (partially) mediated by chromatin interactions.

### Alteration of putative causal genes in the 11p11.2 locus affected AD-related molecular phenotypes

A total of 15 causal genes were predicted to be regulated by functional variants in the 11p11.2 locus and had expressional alterations at the early stage of AD (**Supplementary Table S6**). Moreover, expression profiles from four postmortem brain regions of AD patients and controls generated by our previous study ^32^ showed that most of these genes were also differentially expressed (*P*<0.05) at the late stage of AD (**Fig. 4A**). In order to investigate whether expression alterations of these genes indeed participated in the development of AD, we knocked down and overexpressed four causal genes (i.e. *MADD*, *PSMC3*, *MTCH2*, and *FNBP4*) that were affected by the greatest number of functional variants and had a relatively high expression in the hippocampus (**Fig. 3A**) in U251-APP cells. We measured the levels of the key AD molecules Aβ1-42, Aβ1-40 and phosphorylated tau (pTau-396) by ELISA. Aβ1-40 was undetectable in U251-APP cells and was not included in the subsequent analysis. Knockdown of all four genes led to a significant increase of the Aβ1-42 level in the culture supernatant and the pTau-396 level in the cell lysate, whereas overexpression had opposite effects (**Fig. 4B**). These results indicated the active involvement of multiple causal genes for AD in the 11p11.2 locus.

**Figure 4.**
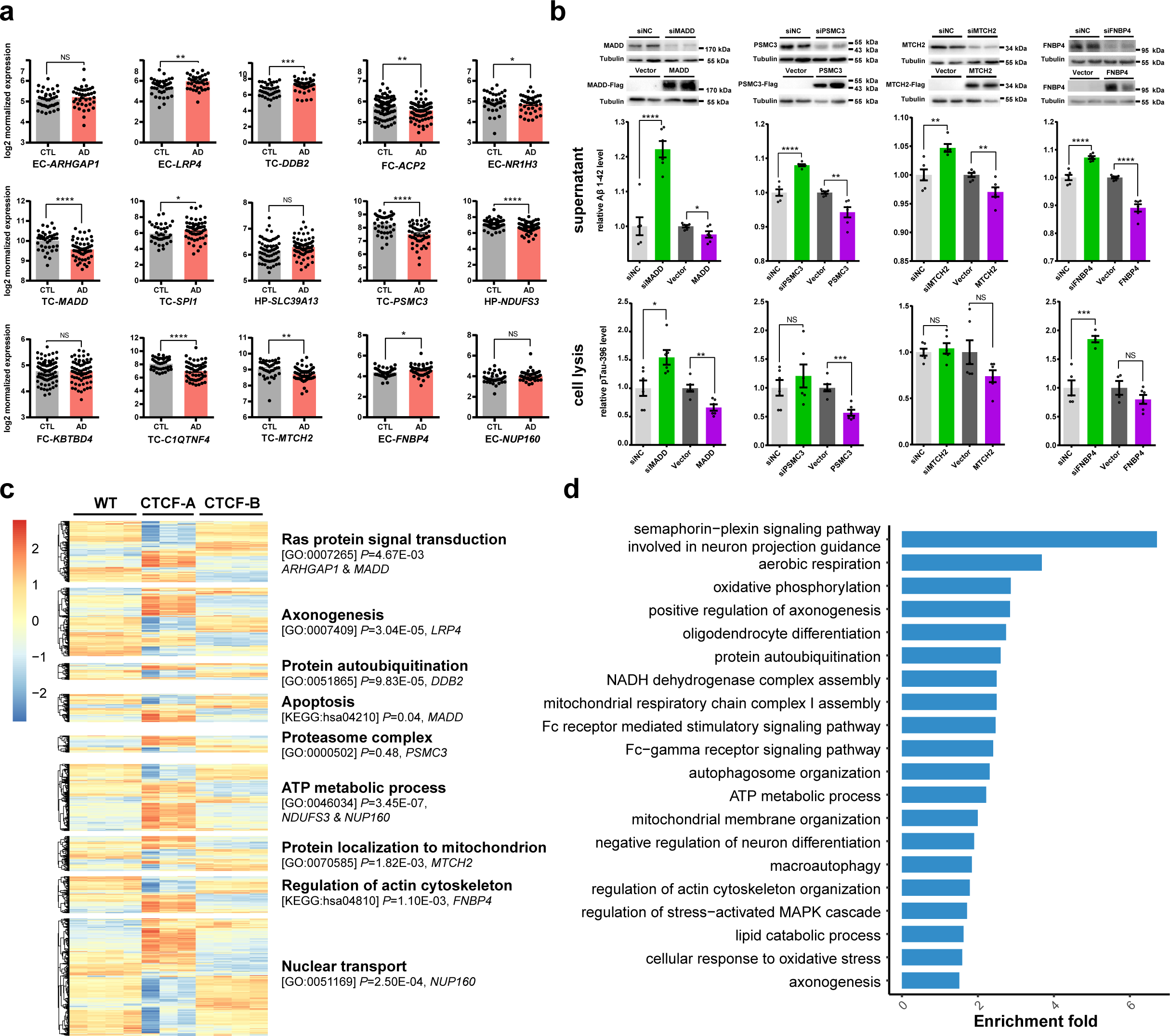
Expression changes of causal genes in the 11p11.2 locus affected AD-related molecular phenotypes and AD-related pathways. **a. Differential expression of putative causal genes in postmortem brain tissues of AD patients and controls.** CTL: controls; AD: AD patients; EC: entorhinal cortex; HP: hippocampus; TC: temporal cortex; FC: frontal cortex. **b. Knockdown or overexpression of four causal genes in the 11p11.2 locus affected Aβ1-42 level in culture supernatant and pTau-396 level in cell lysate of U251-APP cells.** Shown results were a representative experiment of three independent assays with similar results. Values are presented as mean ± SD (n = 6 replicates). **c. Expression changes of causal-gene-relevant pathways disrupted by knocking out CTCF-binding sites.** The enrichment of differentially expressed genes in target pathways in CTCF-A (n = 3 cell clones) and CTCF-B (n = 4 cell clones) cells compared with wild-type cells (WT, n = 3 cell clones) were tested by using the Fisher’s exact test. The corresponding *P*-values (*P*) were listed below each pathway. **d.** Differentially expressed genes in CTCF-A and CTCF-B compared with wild-type cells (WT) were significantly enriched in AD-related pathways (FDR<0.05). *, *P*<0.05; **, *P*<0.01; ***, *P*<0.001; ****, *P*<0.0001; NS, not significant, *P*>0.05; two-sided student’s *t* test.

### Causal genes in the 11p11.2 locus synergistically contributed to AD pathogenesis by affecting multiple AD-related pathways

The causal genes in the 11p11.2 locus might be synergistically regulated by functional variants via chromatin interactions, which possibly made them co-regulated as a whole (**Fig. 3C and Fig. 3D**). In fact, expression levels of these causal genes were predicted to be simultaneously altered by functional variants in the 11p11.2 locus as revealed by the eQTL data (**Fig. 3A**). In order to explore how the concurrent expressional changes of causal genes in the 11p11.2 locus contributed to AD development, we conducted RNA sequencing (RNA-seq) of the cells with knockout of CTCF binding sites in this locus (CTCF-A cell and CTCF-B cell). We found the expression changes of these causal genes could partially mimic the effects induced by functional variants in the 11p11.2 locus (**Fig. 3D**). Genes with an opposite direction of differential expression (FDR<0.05) in CTCF-A and CTCF-B cells were regarded as reliable downstream genes of the 11p11.2 locus (**Supplementary Fig. S15**). These differentially expressed genes were significantly enriched in pathways related to reported functions (**Supplementary Table S7**) of causal genes in the 11p11.2 locus (**Fig. 4C,** Fisher’s exact test *P*<0.05), and most of these pathways were reported to be involved in the etiology of AD ^32, 34–36, 45^. Several other AD-related pathways, such as neuron differentiation, macro-autophagy, stress-activated MAPK cascade, and lipid catabolic process ^32, 46–48^ were also enriched (**Fig. 4D**, **Supplementary Table S8,** FDR<0.05).

### Co-existence of multiple putative causal genes underlying GWAS signals of complex traits

The three LBs in the 11p11.2 locus were found to be implicated in several other complex traits besides AD (**Fig. 5A, Supplementary Table S9**) ^49, 50^. In order to see if the 11p11.2 locus contributed to these traits in a similar mechanism to that of AD, we examined the gene expression alterations contributed by genetic variants in the 11p11.2 locus by integrating eQTL data from trait-related tissues and corresponding GWAS data from public sources ^51–54^ using the MetaXcan algorithm ^51^. Indeed, multiple genes were predicted to be differentially expressed for each trait (**Fig. 5A**, *P*<0.001). Consistently, multiple causal genes identified by the above functional annotation including *ARHGAP1*, *NR1H3*, *NDUFS3*, *C1QTNF4*, *MTCH2*, and *FNBP4* were predicted to be differentially expressed in brain tissues using AD GWAS data (**Fig. 5A**, *P*<0.001). The co-existence pattern of multiple causal genes in a single GWAS signal of complex traits was then further verified by investigating gene expression changes in two datasets related to these traits. The first dataset was composed of expression profiles of pulmonary arterial hypertension patients (PAH) and controls (CTL) ^55^, and 10 eQTL genes (GTEx, lung, *P*<0.001) of the 11p11.2 locus were differentially expressed in lung tissues of PAH compared to controls (**Fig. 5B**, *P*<0.05). Another dataset was a comparative transcriptome analysis of mouse multipotent hematopoietic stem/progenitor cells and myeloid committed precursors ^56^, in which 4 eQTL (GTEx, whole blood, *P*<0.001) genes were identified to be differentially expressed during the stem cell commitment (**Fig. 5C**, *P*<0.05). These results provided additional evidence for the involvement of multiple causal genes in the 11p11.2 locus in modulating complex traits.

**Figure 5.**
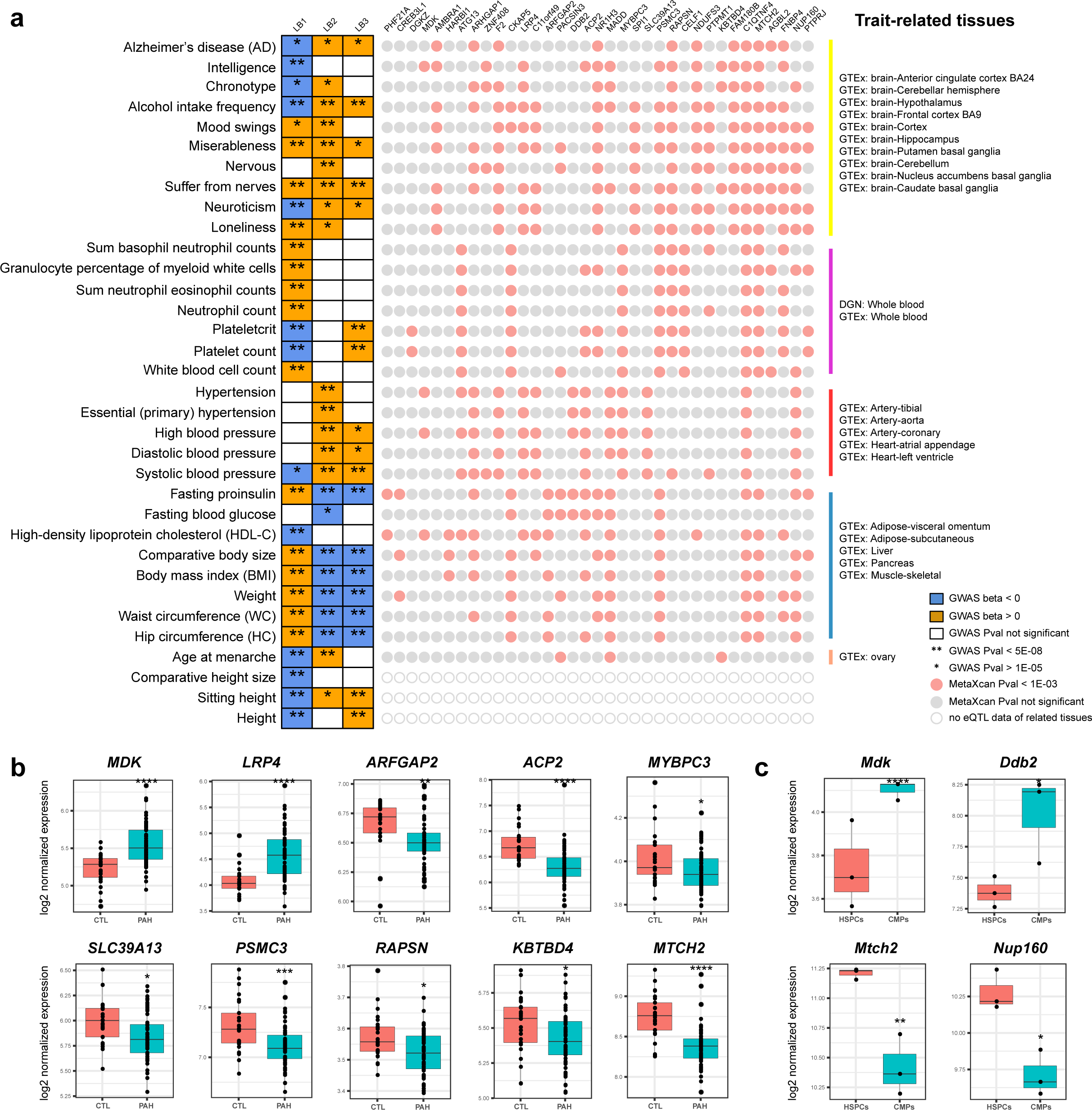
Multiple putative causal genes in the 11p11.2 locus were inferred from GWAS of other complex traits. **a. Predicted differential expression genes caused by genetic variants in the 11p11.2 locus for AD and other complex traits.** LB: long haplotype in the 11p11.2 locus. **b. Differentially expressed eQTL genes (GTEx lung, *P*<0.001) in lung tissues from pulmonary arterial hypertension patients compared with controls (*P*<0.05).** CTL: control; PAH: patients with pulmonary arterial hypertension. **c. Differentially expressed eQTL genes (GTEx whole blood, *P*<0.001) during the hematopoietic stem/progenitor cells differentiation toward myeloid commitment (P<0.05).** HSPCs: hematopoietic stem/progenitor cells; CMPs: common myeloid progenitors.

Collectively, our *in silico* and *in vitro*data provided convincing evidence for the co-existence of multiple functional variants and causal genes within one GWAS locus. For the 11p11.2 locus in AD, we found that one functional variant located in a regulatory element could affect expression levels of multiple proximal and distal genes via chromatin interactions, whereas multiple functional variants in a single locus had a synergistic effect on regulating gene expression in this locus. Alteration of these causal genes influenced the AD-related pathways, which finally contributed to the genetic susceptibility of AD (**Fig. 6**). This *multiple-causal-gene-cluster model* might be a novel mechanism applicable to other GWAS loci of complex traits, and indicated the complex process implicated in the genetic susceptibility of complex traits.

**Figure 6.**
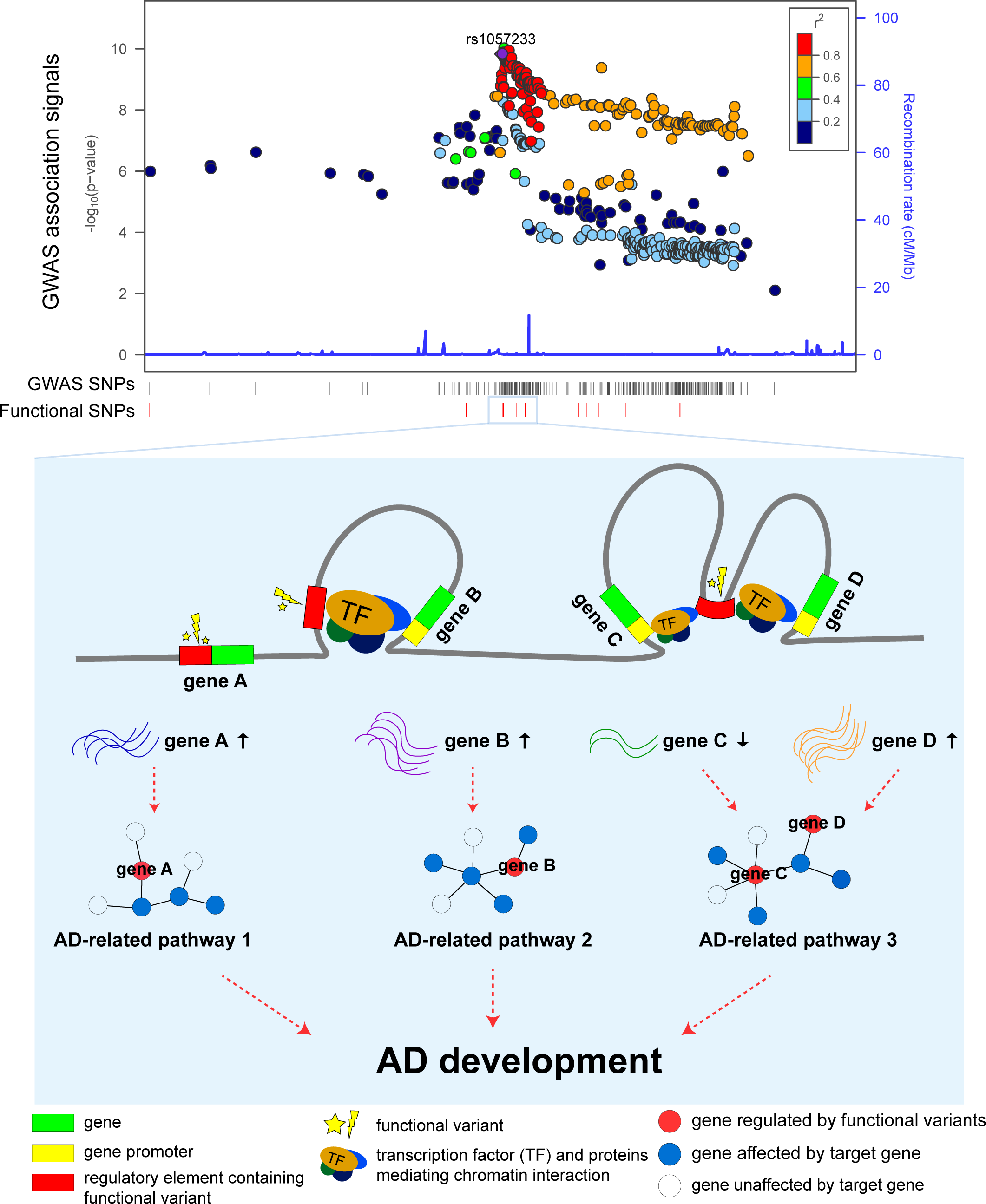
The multiple-causal-gene-cluster model. Multiple functional variants co-exist in a single GWAS locus and regulate expression of a cluster of causal genes via a variety of regulation models. Co-dysregulation of multiple causal genes further disturb gene expression of different pathways related to AD pathogenesis and finally contribute to the development of AD (and other complex traits).

## Discussion

With the remarkable advance of recent AD GWASs, dozens of genetic loci showing robust associations with AD risk have been identified ^4–9^. Whereas, the decipherment of real responsible variants and genes within these loci as well as related biological mechanisms is still a conundrum. In this study, we conducted a systematic functional genomic study of AD risk loci by integrating multiple levels of omics data, and identified 304 functional variants and 166 causal genes for AD. We found that there were multiple functional variants and causal genes for most of AD GWAS loci. Those putatively causal variants contributed to AD risk through regulating a functionally synergetic gene cluster, as shown by cellular experiments of the 11p11.2 (CELF1/SPI1) locus here. Further analyses showed that the *multiple-causal-gene-cluster* pattern is common for GWAS loci of AD and other complex traits. This model might be a general mechanism for interpreting GWAS and post-GWAS results.

Great efforts were made to annotate and interpret AD GWAS signals, especially by the integration of eQTL and epigenetic annotations ^5, 6, 11, 17–19, 57^. These studies have moved a critical step from loci to variants and genes, however, complex LD structure, limited types or sample size of annotation data, and lack data from AD-specific cell types or tissues made the fine-mapping somewhat incomplete and elusive for follow-up experimental validation. In this study, well-curated multi-omics data with relatively large sample size from AD-related tissues or cells were used for annotation. AD-related SNPs in loci with genome-wide or sub-threshold significance from three large-scale AD GWASs ^5, 6, 8^ were integrated with these annotation data, which allowed for a more systematic and unbiased screening of AD risk loci. As a result, we prioritized additional putative functional variants and causal genes for most of AD GWAS loci, and observed a multiple-causal-gene-cluster pattern in many loci. We also provide a reliable list of potentially functional variants and causal genes for AD (available at www.alzdata.org), which may act as a promising resource for further mechanistic and therapeutic studies. We further validated the reliability of the prioritized functional variants and causal genes and investigated the underlying regulatory mechanisms by cellular assays. Importantly, we have proposed a *multiple-causal-gene-cluster model* to interpret the GWAS signals, providing a framework towards a mechanistic interpretation of how statistically-associated SNPs affect biological processes of complex traits.

The 11p11.2 locus was one of the most complicated GWAS risk locus for AD, with complex linkage structure and eQTL signals. The observation that multiple causal genes might be co-existed in this locus was noticed in several studies ^5, 17, 57^. Karch et al. identified several genes with eQTL signals of the 11p11.2 locus and were also differentially expressed in AD brains, and they speculated that genes within this region might act cooperatively to modify AD risk and there was a key regulator within this region influencing the expression of many other genes ^17^. Subsequent study focusing on this locus suggested that the downregulation of transcription factor SPI1 (PU.1) caused by the protective haplotype in the 11p11.2 locus was responsible for a lower AD risk, and SPI1 might be a key regulator for AD (Huang et al., 2017). Recently, Novikova et al. found that several genes shared myeloid active enhancers that contain AD risk alleles in the 11p11.2 locus. They speculated that either multiple causal genes or one single true causal gene with several its co-expressed risk-neutral genes were present in this locus ^57^. Our data showed that the co-regulation of a *multiple-functional-gene-cluster*, rather than a key regulator, accounted for the association signal of the 11p11.2 locus. Indeed, other genes in this region, e.g. *MADD* ^58^ and *MTCH2* ^41^, were proved to be involved in AD-related molecular changes by *in vivo* data. Additionally, proteasome component gene *PSMC3* and nuclear pore complex gene *NUP160* were reported as potential causal genes in the 11p11.2 locus, whereas genes encoding subunits of these components were also prioritized as causal genes for other GWAS loci in our study, including *PSMG3* ^59^, *NUP85* and *NUP88* ^60^. All these observations proved the active involvement of these genes/pathways in AD according to the omnigenic theory ^61^. Moreover, our cellular assays provided direct evidence showing the co-existence of several functional variants and multiple causal genes, and elucidated the underlying mechanism that these variants and genes synergistically contributed to AD risk.

The *multiple-causal-gene-cluster model* was not restricted to the 11p11.2 locus, but was also applicable to other AD GWAS loci. Our results showed that most of GWAS loci harbored more than one functional variant and causal gene, especially for the 7q22.1, 16p11.2, and 19q13.31-13.32 locus, which had a comparatively complex LD and gene regulation as that of the 11p11.2 locus. The *APOE Ɛ4* allele was regarded to be the main contributor of the 19q13.31-13.32 locus for a long time, however, plenty of genome-significant variants resided in this locus were not in LD with SNPs defining the *APOE* types (rs429358 and rs7921, *R*^2^<0.2) (**Supplementary Fig S13**), suggesting potentially independent effect. In addition, recent studies reported several AD susceptible coding and non-coding variants except for *APOE Ɛ4* ^62–64^, and a total of eight genes (*APOE*, *TOMM40*, *PVR*, *BCAM*, *APOC1*, *APOC4*, *CLPTM1*, and *IGSF23*) were implicated by functional interpretation strategies in the Jansen study ^6^. Consistently, we identified 63 functional variants and 19 causal genes for this locus in the current analyses. What was similar to the 11p11.2 locus is that causal genes in this locus were implicated in different pathways, including lipid metabolism (*APOE*, *APOC1* and *NKPD1* ^45, 65, 66^), immune process (*PVR*, *PVRL2*, *CEACAM19*, and *BCL3* ^67–69^), and DNA repair (*ERCC1* and *ERCC2* ^70, 71^). For the 7q22.1 locus, Kikuchi et al. ^18^ found an enhancer variant connected many eQTL genes via the CTCF-mediated chromatin loops, which was also exemplified in our results. Besides AD, the *multiple-causal-gene-cluster model* might also exist in other complex traits as revealed by this study and others ^14, 15^. Therefore, the co-existence of multiple functional variants and causal genes within one GWAS locus might be common to complex traits. This *multiple-causal-gene-cluster model* might act as a potential link filling the gap between functional variants and biological mechanisms.

In conclusion, we systematically characterized the regulatory mechanisms of AD risk loci, validated the regulatory effects with cellular experiments, and translated the genetic associations to specific genes. We illustrated how the identified functional SNPs could confer risk of AD by regulating multiple genes in the 11p11.2 locus. We proposed a *multiple-causal-gene-cluster model* for interpreting GWAS results and moved one step beyond association towards mechanism by assessing the effects of multiple SNPs within one locus, adding novel insights into the genetic mechanisms of AD and other complex traits. Our results offered an updated perspective regarding candidate genes mapping for further functional characterization and drug development. Further studies, such as high-throughput validation of functional variants, cell-type-specific regulation effect of functional variants/genes, and *in vivo* assays of candidate genes, are needed to confirm the involvement of the multiple causal genes in AD and to elucidate the synergetic mechanisms of gene clusters in complex traits.

## Online content

Any methods, additional references, Nature Research reporting summaries, source data, supplementary information, details of author contributions and competing interests; and statements of data and code availability online.

## Methods

### GWAS summary datasets and candidate variants for AD

The genetic association results from three large-cohort GWASs for AD were used in this study ^5, 6, 8^. The Lambert study ^8^ and the Kunkle study ^5^ were conducted by the International Genomics of Alzheimer’s Project (IGAP). Summary-level statistics from stage 1 of the Lambert study ^8^ and the Kunkle study ^5^ were used for filtrating candidate variants ^5, 8^. The Lambert dataset included 17,008 AD patients and 37,154 controls (Nsnps=7,055,881) ^8^. The Kunkle dataset added 17 new datasets to the Lambert study, and resulted in 21,982 AD patients and 41,944 controls (Nsnps= 36,648,992) (Kunkle et al., 2019). The Jansen study ^6^ was based on clinical diagnosed AD and AD-by-proxy (individuals with one or two parents with AD) samples. Phase 3 of the Jansen study was a meta-analysis of the Lambert study stage 1 data (N=54,162) ^8^, AD GWAS from the Psychiatric Genomics Consortium (N=17,477), whole-exome sequencing (WES) data from the Alzheimer’s Disease Sequencing Project (ADSP, n=7506), and GWAS of AD-by-proxy subjects and controls from the UK Biobank (N= 376,113). A total of 13,367,299 variants were genotyped or imputed in this dataset (Jansen et al., 2019). All participants in these studies were of European origins.

Common variants (minor allele frequency [MAF]>0.01) with GWAS *P*-value reached genome-wide (*P*<5x10^-8^) or sub-threshold significance (P<1x10^-5^) and those in linkage with these variants (LD, *R^2^*>0.8) were defined as candidate risk variants for AD. Genotype data from the 1000 Genomes project phase 3 (European population, EUR) were used to compute LD among variants ^72^. SNPs in the MHC region (chr6: 28500000-33500000) were excluded from analysis because of its complex linkage structure. The resulting variants from all three studies were combined to achieve the final candidate variant set (Nsnps=7884) and were subjected to functional annotation.

### Functional annotation of AD candidate variants

Multi-layers of omics data were used to prioritize functional variants for AD from the above AD candidate variant dataset. In order to identify variants located in active regulatory elements and with abilities affecting transcriptional events, data of histone modification, open chromatin, transcription factor (TF) binding, and expressional quantitative trait loci (eQTL) were used for functional annotation. Briefly, histone modification data for promoters (H3K4me3 and H3K9ac) ^23^ and enhancers (H3K4me1 and H3K27ac) ^24^ were downloaded for 8 brain regions (layer of hippocampus, temporal lobe, angular gyrus, caudate nucleus, cingulate gyrus, middle frontal area 46, substantia nigra, and embryonic brain) and 6 neural cells (astrocyte, bipolar neuron, neural cell, neural stem progenitor cell, neuron, and radial glial cell) from the Encyclopedia of DNA Elements (ENCODE) (https://www.encodeproject.org) ^21, 22^. Histone modification data for monocyte (CD14-positive monocyte) were also included for analysis because of its active role in AD ^40^. ENCODE accessions for these datasets were listed in **Supplementary Table S1**. Bed files for these histone modification markers of these tissues or cell types were used to annotate candidate variants. A false discovery rate (FDR) < 0.001 was used to filter peaks. A variant was considered to locate in a promoter or enhancer only if it located in promoter/enhancer peaks in at least 2 brain tissues or neural cells.

Assay for transposase accessible chromatin using sequencing (ATAC-seq) data of neural tissues or cells were used to identify variants located in transcriptional active regulatory regions. ATAC peaks of iPSC-induced excitatory neurons, iPSC-derived hippocampal DG-like neurons, and primary fetal astrocytes were downloaded from Gene Expression Omnibus (GEO, https://www.ncbi.nlm.nih.gov/geo/) with accession number GSE113483 ^25^. ATAC peaks of neuron and glia cell from 14 brain regions were downloaded from the Brain Open Chromatin Atlas (BOCA, http://icahn.mssm.edu/boca) ^26^. ATAC of monocyte was downloaded from the GEO database with accession number GSE87218 ^27^. Peaks with FDR < 0.001 were used to intersect with the candidate variants.

In order to identify variants whose different alleles might affect the binding affinity of transcription factors, chromatin immunoprecipitation sequencing (Chip-seq) data of over 600 TFs from the ENCODE were used to identify variants located in transcription factor binding sites (TFBS) ^21, 22^. ENCODE accessions for these datasets were listed in **Supplementary Table S2**. FDR<0.05 was used to filter peaks from bed files. DNA sequences of the top 1000 peaks (ranked by peak height) for each transcription factor (TF) were subjected to MEME to predict the DNA binding motifs (position weight matrix, PWM) (-mod zoops -nmotifs 3 -minw 6 -maxw 30). Top 3 PWMs with the smallest *E*-values for each TF were subjected to R package atSNP ^28^ to predict whether different alleles of variants resided in the TFBS could affect binding affinities of this TF. A variant was considered to disrupt the TF binding affinity if DNA sequence with reference allele (Pval_ref<0.05) or alternative allele (Pval_alt<0.05) of this variant was able to bind with the target TF, and their binding affinities were significantly different (Pvalue_rank<0.05).

The eQTL data of brain tissues and monocytes were used to verify whether different alleles of a variant were associated with different expression levels of a gene. eQTL data for brain tissues included the eMeta dataset ^7^ and the PsychENCODE dataset ^29^. The eMeta dataset was a meta-analysis ^7^ of brain tissues from the Genotype-Tissue Expression (GTEx) project ^52, 53^, the CommonMind Consortium (CMC) ^73^, and the Religious Orders Study and Memory and Aging Project (ROSMAP) ^74^ with an effective sample size of 1194. The PsychENCODE dataset had a comparable sample size with the eMeta dataset, which included 1387 samples from the prefrontal cortex ^29^. eQTL data of these two datasets were downloaded from the SMR website (http://cnsgenomics.com/software/smr/#eQTLsummarydata) and extracted by using the SMR software ^75^. The eQTL dataset for monocytes were from Raj et al. ^31^ and Kim-Hellmuth et al. ^30^ with sample size of 461 and 134, respectively. Only cis-eQTL results (SNPs within ±1 megabases [Mb] of the target gene) were used in this analysis. A *P*-value < 0.001 was considered as statistically significant. The significant cis-eQTL gene of the target SNP was defined as eGene of the SNP.

### Differential expression of target genes at bulk-tissue level and single-cell level

Normalized expression and differential expression results of AD patients and controls at the bulk-tissue level in entorhinal cortex (EC), hippocampus (HP), temporal cortex (TC), and frontal cortex (FC) for target genes were obtained from our previous study (AlzData: www.alzdata.org) ^32^. Raw counts of single-cell RNA sequencing data of AD patients and controls were applied from the Synapse database (https://www.synapse.org/#!Synapse:syn18485175) and normalized by using the Seurat R package running the default parameters ^33, 76^. Differential expression results of target genes between AD patients and controls were from the original study ^33^. Differentially expressed gene was defined when the FDR-corrected *P*-value < 0.05 in the two-sided Wilcoxon-rank sum test reported by the original study ^33^.

### Detection of long linkage blocks (LBs) for the 11p11.2 locus

The summary-level statistics from the Lambert study ^8^ for the 11p11.2 locus were clumped using Plink v1.9 (www.cog-genomics.org/plink/1.9/) ^77^ by using genotyping data from the 1000 Genomes Project phase 3 (EUR population) as reference (--clump-p1 1E-5 --clump-kb 1000 --clump-r2 0.8) ^72^. Long linkage blocks (LBs) were defined as linkage blocks constituted by over 50 SNPs.

### Vector construction and dual-luciferase reporter assays for candidate functional variants

The DNA fragment of regulatory region containing the target SNP was amplified from in house human DNA samples ^64, 78^ by using primers listed in **Supplementary Table S10** and was inserted into the pGL3-basic (for promoter assays) or pGL3-promoter (for enhancer assays) luciferase reporter vector. For SNPs located in promoter regions, including rs11039159, rs7947450, rs2280231, rs35624992, and rs11039376, the DNA fragments containing these SNPs were inserted into PGL3-basic vector. For SNPs located in enhancers, including rs11039149, rs11039225, rs2293577, rs71475921, and rs7120548, the DNA fragments were inserted into PGL3-promoter. PCR-mediated point mutation was used to generate DNA fragments containing the respective alternative alleles of target SNPs by using primers in **Supplementary Table S10**. All sequences of the inserted DNA fragments were verified by Sanger sequencing.

The HEK293T cells were used to perform the luciferase reporter assay. Cells were introduced from Kunming Cell Bank, Kunming Institute of Zoology, and were cultured in Dulbecco’s modified Eagle’s medium (DMEM; Thermo Fishers) supplemented with 10% fetal bovine serum (FBS, Thermo Fishers) at 37 °C in 5% CO2. The constructed vector (250 ng per well for 48-well plate) and the internal control vector TK (25 ng per well for 48-well plate) were co-transfected into the cells by using the X-tremeGene HP DNA transfection reagent (ROCHE, 6366236001). The transfected HEK293T cells were harvested in 65 μL passive lysis buffer (Promega, USA) at 24 h. Luminoskan Ascent instrument (Thermo Scientific) was used to measure the luciferase activity with the Dual -Luciferase Reporter Assay System (Promega, E1910). Six replicates were used for each vector and all experiments were conducted in triplicates. Two-tailed Student’s *t* test was performed to quantify the statistical difference between the two groups by using GraphPad Prism software (GraphPad Software, La Jolla, CA, USA). A *P*-value < 0.05 was considered as statistically significant.

### Base-editing of target functional variants

The CRISPR-based genome-editing approach base-editing was conducted to directly generate precise point mutations into cellular DNA for typical functional variants. Guide RNAs (gRNAs) targeting to the genomic region of rs2280231 and rs2297533 were designed (**Supplementary Table S10**), and were sub-cloned into the pGL3-U6-sgRNA-PGK-puromycin (Addgene plasmid # 51133) plasmid. The HEK293T cells were cultured in DMEM medium (Thermo Fishers) supplemented with 10% FBS (Thermo Fishers). Cells were seeded in a 6-well plate with a density of 5 × 10^5^ per well 12 h before transfection. Constructs containing different sgRNAs (500 ng) were co-transfected with pCMV-ABE7.10 (2000 ng, Addgene plasmid # 102919) into cells by using Lipofectamine^TM^ 3000 (Thermo Fishers) according to the manufacture’s manual. The pCMV-ABE7.10 ^79^ was a gift from Prof. Xingxu Huang. Twenty-four hours after transfection, medium was changed daily with fresh medium supplemented with 2 μg/mL puromycin, and cells were selected by puromycin for 5 days. Single cells resistant to puromycin were seeded in 96-well plate and were cultured for 2-3 weeks to obtain single cell clones. Genomic DNA of each clone was extracted by using the AxyPrep Genomic DNA Miniprep Kit (Axygen, AP-MN-BL-GDNA-250). The target region was amplified and sequenced to confirm successful base-editing of rs2280231 and rs2297533.

### Detection of topologically-associated domains (TADs) in the 11p11.2 locus

Promoter capture Hi-C (pcHi-C) data for induced pluripotent stem cell (iPSC)-derived hippocampal dentate gyrus (DG)-like neurons were used to detect TADs in the 11p11.2 locus ^25^. CHiCAGO interaction data in IBED format were downloaded from the Gene expression Omnibus (https://www.ncbi.nlm.nih.gov/geo/) with accession number GSE113481. The ±1Mb of the 11p11.2 locus (chr11: 46000000-49000000) were extracted from the interaction file. R package SpectralTAD was used to detect TADs by using a resolution of 10 kilobases (kb) ^80^. The interaction heatmap was plotted by using the R package HiTC ^81^. pcHi-C interactions from iPSC-derived hippocampal DE-like neurons and astrocytes were applied to visualize chromatin interactions among loci in the 11p11.2 locus ^25^. More details regarding the data information can be found in the original paper ^25^.

### Deletion of the CTCF-binding sites using the CRISPR/Cas9

Two guide RNAs (gRNAs) targeting to the upstream and downstream region of the CTCF-binding sites (CTCF-A and CTCF-B) were designed according to the PAM sequence “NGG” in this region (**Supplementary Table S10**). The two gRNAs were respectively sub-cloned into the pGL3-U6-sgRNA-PGK-puromycin (Addgene plasmid # 51133) plasmid and PX330-mcherry (Addgene plasmid #98750) plasmid, and were co-transfected into U251-APP cells using Lipofectamine™ 3000 (Invitrogen L3000008). The PX 330-mcherry ^82^ was a gift from Prof. Jinsong Li, and the pGL3-U6-sgRNA-PGK-puromycin ^83^ was a gift from Prof. Xingxu Huang. U251-APP cells were cultured in Roswell Park Memorial Institute 1640 Medium (Thermo Fishers) supplemented with 10% FBS (Thermo Fishers).

Forty-eight hours after transfection, single cells were picked from puromycin-resistant cells with strong mcherry fluorescence signals and were seeded and cultured in 96-well plate for 3-4 weeks to obtain single cell clones. Genomic DNA of each clone was extracted by using the AxyPrep Genomic DNA Miniprep Kit (Axygen, AP-MN-BL-GDNA-250). The target region was amplified and sequenced to confirm successful deletion of the targeted CTCF-binding site.

Potential off-target sites in cells edited by the CRISPR/Cas9 were predicted using the Breaking-Cas (https://bioinfogp.cnb.csic.es/tools/breakingcas/) ^84^. Off-target sites with no more than 3 mismatches with corresponding primers used for CRISPR/Cas9 with score<1 by the Breaking-Cas were checked (**Supplementary Table S11**). DNA regions containing off-target sites were amplified by PCR and validated using Sanger sequencing.

### Knockdown or overexpression of the causal genes in U251-APP cells

The U251-APP cells were used for testing the effect on AD-related molecular phenotypes caused by knockdown or overexpression candidate causal genes. U251-APP cells were cultured in Roswell Park Memorial Institute 1640 Medium (Thermo Fishers) supplemented with 10% FBS (Thermo Fishers) at 37 °C in a humidified atmosphere incubator with 5% CO2, as described in our previous studies ^32, 43, 44^. Cells (2 × 10^5^ per well) were seeded in 6-well plates to grow to 50% confluence for transfection. siRNA of target gene (20 nM) or control siRNA (siNC) was dissolved in Opti-MEM medium, and was then mixed with 5 μL Lipofectamine^TM^ 3000 (Invitrogen, L3000008) to achieve a final volume of 100 μL. The siRNA-lipofectamine mixture was incubated at room temperature for 10 min, and then was added to each well. After an incubation for 6 h, medium was removed and fresh medium was added to each well (2 mL) for growth.

Overexpression plasmids were respectively transfected into U251-APP cells by using the X-tremeGene HP DNA transfection reagent (ROCHE, 6366236001). Cells grown in 6-well plate were transfected by overexpression plasmid of the target gene or its corresponding control plasmid (each 2 μg) together with HP transfection reagent (4 μL) dissolved in 200 μL Opti-MEM medium. The mixture was incubated at room temperature for 10 min, then was added to each well. Culture supernatant in each well transfected with siRNA or overexpression vector was replaced with equal volume of new growth medium after 24 h, and 1 μg/mL doxycycline (Sigma; D9891) was added to induce APP expression. Cells were harvested at 72 h after transfection.

### Western blot

Cells were lysed with protein lysis buffer (Beyotime, P0013) on ice and centrifuged at 12 000×g at 4 ^◦^C for 10 min to remove cell debris. Protein concentration was determined using a BCA Protein Assay Kit (Beyotime, P0012). A total of 20 μg protein was separated with 12% (vol/vol) sodium dodecyl sulphate (SDS)-polyacrylamide gel and electrophoretically transferred onto a polyvinylidene difluoride membrane (Bio-Rad, L1620177). The membrane was soaked with 5% (w/v) skim milk for 2 h at room temperature, and then was incubated with the primary antibodies against MADD (1:1000; abcam, ab134117), MTCH2 (1:1000; absin, abs143485), PSMC3 (1:1000; abcam, ab171969), FNBP4 (1:1000; ABclonal, A13804), Flag (1:5000; Abmart, M20008), and Tubulin (1:20000; EnoGene, E1C601) overnight at 4 ^°^C, respectively. After three washes with Tris-buffered saline with 0.1% Tween detergent (each 5 min), the membranes were incubated for 1 h with anti-mouse or anti-rabbit secondary antibodies (1:10000, KPL, USA) at room temperature. The membrane was visualized using enhanced chemiluminescence reagents (Millipore, WBKLS0500).

### ELISA analysis of Aβ1-42, Aβ1-40, and pTau-396

The levels of Aβ1-42 (Elabscience, E-EL-H0543c) and Aβ1-40 (Elabscience, E-EL-H0542c) in culture supernatant, and pTau-396 (Elabscience, E-EL-H5314c) in cell lysate of U251-APP cells with different conditions were measured by using commercial enzyme-linked immunosorbent assay (ELISA) kit. A total of 100 μL culture supernatant of cells or cell lysate was used to perform the ELISA experiment according to the manufacturer’s instructions.

### RNA extraction and Quantitative real-time PCR (RT-qPCR)

Total RNA was extracted by using the RNAeasy kit (TianGen) according to the manufacturer’s instructions. The A260/A280 ratio of total RNA was measured on a NanoDrop biophotometer (Thermo Fisher Scientific). Only samples with A260/A280 ratio of 1.8–2.0 were used for subsequent experiments. Quality and integrity of RNA samples were evaluated based on the 28S and 18S rRNA bands on a 1% agarose gel. Around 1 μg total RNA was used to synthesize cDNA by using oligo-dT18 primer and Moloney murine leukemia virus reverse transcriptase (M1701, Promega). The RT-qPCR was performed using iTaq Universal SYBR Green Supermix (172-5125, Bio-Rad Laboratories) supplemented with gene-specific primer pairs (**Supplementary Table S10**) on a CFX Connect Real-Time PCR Detection System (Bio-Rad Laboratories). The *ACTB* transcript was used for the normalization of the target gene.

### RNA sequencing (RNA-seq) and data analysis

About 1.5 μg total RNA per sample was used to prepare library for RNA-seq. Sequencing libraries were generated using the NEBNextUltraTM RNA Library Prep kit for Illumina (NEB, USA) following the manufacturer’s recommendation and index codes were added to attribute sequence to each sample. The final library was sequenced on an IlluminaHiseq 4000 platform, and 150 bp paired-ends reads were generated. Raw RNA-seq reads were trimmed to remove sequencing adapters and low-quality reads by using the Trimmomatic (version 0.33) ^85^. The clean reads were then aligned to the reference genome (GRCh38.p7) using HISTA2 ^86^. FeatureCounts was used to generate counts for known human genes ^87^. Gene-level differential expression analyses were performed using DESeq2 ^88^.

### Exploration of genetic associations of the 11p11.2 locus with other complex traits

The GWAS results from the GWAS Catalog (https://www.ebi.ac.uk/gwas/) ^50^ and a total of 600 GWASs from the gwasATLAS (https://atlas.ctglab.nl/) ^49^ were used to explore the potential implication of the 11p11.2 locus in other complex traits. Those traits with over 50% SNPs in one of the three blocks of the 11p11.2 locus reaching sub-threshold significance (GWAS *P*<1x10^-5^) was considered as 11p11.2 locus-related. The direction of effect (beta) for each block in each trait was harmonized by defining the same effect allele according to AD GWAS (phase 1 data from the Lambert study ^8^).

### Inferring causal genes for the 11p11.2 locus by integrative analysis

MetaXcan ^51^ was used to map causal genes for the 11p11.2 locus. eQTL data of 48 tissues from the GTEx project ^52, 53^ and whole blood from the Depression Genes and Networks (DGN) ^54^ were used for analysis. The pre-calculated GTEx Version 7 databases and the DGN database for MetaXcan were downloaded from the PredictDB Data Repository (http://predictdb.org/) ^89^. Summary statistic data for the 11p11.2 locus from each GWAS were integrated with the pre-calculated eQTL database from GWAS-related tissues. The statistically significant threshold was set at *P*<0.001.

### Detection of differentially expressed genes located in the 11p11.2 locus in other datasets

Expression profiles for lung tissues from 58 patients with pulmonary arterial hypertension (PAH) and 25 controls were downloaded from GEO (https://www.ncbi.nlm.nih.gov/geo/; accession ID GSE117261) ^55^. Differential expression analysis of PAH patients versus controls was conducted by R package limma ^90^ using gender as covariate. Differential expression results of mouse multipotent hematopoietic stem/progenitor cells versus myeloid committed precursors were reported by the original study ^56^. Detailed eQTL results for the 11p11.2 locus for the lung and whole blood were downloaded from the GTEx Portal (https://www.gtexportal.org/) ^52, 53^.

### Statistical analysis and data visualization

Circular plot for functional annotation results was generated by R package circlize ^91^. Chow-Ruskey diagram for candidate variants and functional variants in each study was plotted by R package Vennerable ^92^. Functional enrichment analysis (Gene ontology, biological process) was performed by R package clusterProfiler ^93^. Heatmap of expression of target gene set was generated by R package pheatmap ^94^. Locuszoom (http://locuszoom.org/) was used to visualize GWAS results ^95^. Functional annotation data for target genomic region was visualized using the WashU epigenome browser (http://epigenomegateway.wustl.edu/) ^96^. LD matrix of target SNPs were generated by Haploview v4.1 ^97^ using genotyping data from the 1000 Genomes Project phase 3 (EUR population) as reference ^72^. Comparisons of relative luciferase activities, mRNA or protein levels between two groups in this study were conducted by the Student’s *t* test using the PRISM software (GraphPad Software, Inc., La Jolla, CA, USA).

### Data availability

Summary statistics of AD GWAS were downloaded from http://web.pasteur-lille.fr/en/recherche/u744/igap/igap_download.php (the Lambert study), https://www.nature.com/articles/s41588-019-0358-2 (the Kunkle study), and https://ctg.cncr.nl/documents/p1651/AD_sumstats_Jansenetal_2019sept.txt.gz (the Jansen study). GWAS results of other traits were downloaded from the GWAS Catalog (https://www.ebi.ac.uk/gwas/) and gwasATLAS (https://atlas.ctglab.nl/). ChIP-seq data of Transcription Factors, H3K4me3, H3K9ac, H3K4me1, and H3K27ac were downloaded from the Encyclopedia of DNA Elements (ENCODE) (https://www.encodeproject.org). ATAC-seq data of iPSC-induced excitatory neurons, iPSC-derived hippocampal DG-like neurons, and primary fetal astrocytes were downloaded from Gene Expression Omnibus (GEO, https://www.ncbi.nlm.nih.gov/geo/) with accession number GSE113483. ATAC peaks of neuron and glia cell were downloaded from the Brain Open Chromatin Atlas (BOCA, http://icahn.mssm.edu/boca). ATAC of monocyte was downloaded from the GEO database with accession number GSE87218. eQTL data were downloaded from the SMR website (http://cnsgenomics.com/software/smr/#eQTLsummarydata) and the GTEx Portal (https://www.gtexportal.org/). Promoter capture Hi-C (pcHi-C) data of iPSC-derived hippocampal dentate gyrus (DG)-like neurons were downloaded from the GEO (https://www.ncbi.nlm.nih.gov/geo/) with accession number GSE113481. Expression profiles for lung tissues were downloaded from GEO with accession ID GSE117261. Raw counts of single-cell RNA sequencing data of AD patients and controls were applied from the Synapse database (https://www.synapse.org/#!Synapse:syn18485175). All detailed annotations for functional variants and causal genes prioritized generated in this study are available at AlzData (www.alzdata.org). RNA-sequencing data of CTCF-binding site knockout cells (CTCF-A and CTCF-B) and controls are available through GEO (accession number will be available before publication).

## Code availability

All custom code used in this work is available at http://www.alzdata.org/download1.php.

## Acknowledgements

The authors thank Ian Logan for language editing and helpful comments to this manuscript. The study was supported by the National Natural Science Foundation of China (31730037 to Y.G.Y., 82022017 and 31970965 to D.F.Z.), the Strategic Priority Research Program (B) of CAS (XDB02020003 to Y.G.Y.), the Project for International Collaboration of the Bureau of International Collaboration, CAS (GJHZ1846 to Y.G.Y.), the Bureau of Frontier Sciences and Education, CAS (QYZDJ-SSW-SMC005 to Y.G.Y.), the CAS “Light of West China” Program and the Youth Innovation Promotion Association of CAS (to D.F.Z.), the Applied Basic Research Foundation of Yunnan Province (to M.X.). We appreciate the ENCODE Consortium and the ENCODE production laboratories for the sharing of epigenetic data. The single-cell level expression data of AD patients and controls used in this study were provided by the Rush Alzheimer’s Disease Center, Rush University Medical Center, Chicago. Data collection was supported through funding by NIA grants P30AG10161, R01AG15819, R01AG17917, R01AG30146, R01AG36836, U01AG32984, U01AG46152, the Illinois Department of Public Health, and the Translational Genomics Research Institute.

## Author Contributions

M.X., Y.G.Y., and D.F.Z. conceived and designed the study. M.X., Y.G.Y., and D.F.Z. compiled the figures and wrote the manuscript with help and input from all authors. Q.L., R.B., M.X., D.F.Z., Y.L., Z.Y., Q.Z., X.L., and C.S. performed experiments. M.X., M.Y., and C.Z. performed the data processing and analysis. X.J.L and M.L. revised the draft. All authors revised the manuscript and approved the publication.

