## Supplementary Figures for "A *multiple-causal-gene-cluster* model underlying GWAS signals of Alzheimer’s disease"

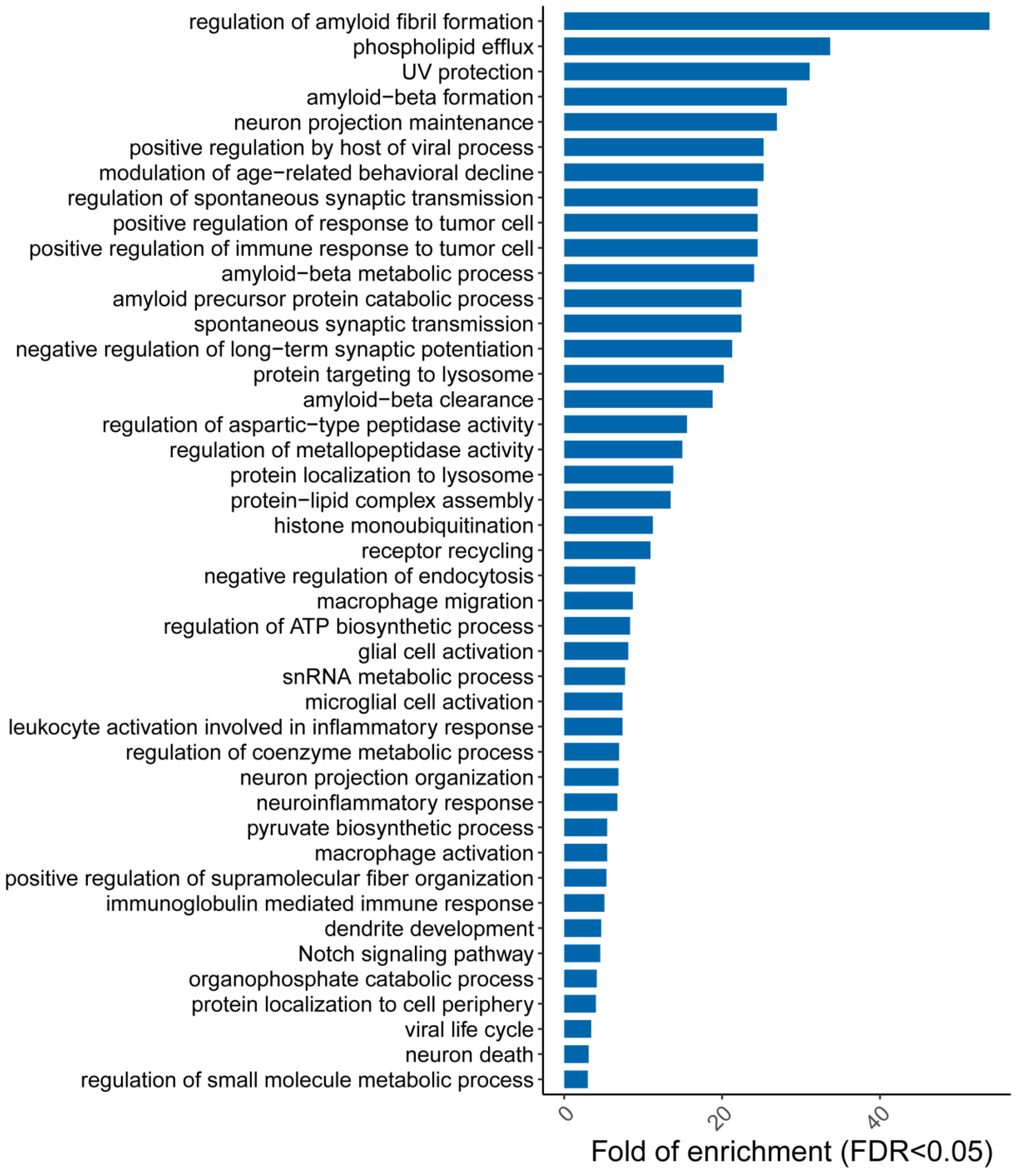


**Supplementary Figure S1**. Enriched pathways (Gene ontology, biological process) of 166 causal genes for Alzheimer's disease (AD), together with known AD pathogenic genes and core regulators (i.e. *APP*, *PSEN1*, *PSEN2*, *TREM2*, *SORL1*, and *TYROBP*) [1-5].

## A.

Functional Variants for AD

Show 10 entries    Search:

| SNP | CHR | POS | A1 | A2 | Gene | Location | IGAP_Beta | IGAP_Pvalue | Kunkle_Beta | Kunkle_Pvalue | Jansen_Beta | Jansen_Pvalue | Histone_mod | ATAC | diff_TF | eQTL |
|---|---|---|---|---|---|---|---|---|---|---|---|---|---|---|---|---|
| rs1004173 | 6 | 47445017 | T | C | CD2AP | upstream | 0.097 | 4.55E-8 | 0.083 | 1.76E-7 | 0.014 | 5.64E-9 | 58 | 9 | 2 | 1 |
| rs10086852 | 8 | 27238115 | G | C | PTK2B | intronic | 0.053 | 6.62E-4 | 0.051 | 4.75E-4 | 0.008 | 1.54E-4 | 9 | 1 | 3 | 1 |
| rs10200967 | 2 | 127841769 | C | T | BIN1 | intronic | -0.087 | 3.05E-6 | -0.097 | 4.31E-9 | -0.015 | 7.61E-10 | 38 | 16 | 2 | 1 |
| rs10212593 | 3 | 155347487 | T | G | PLCH1 | intronic | -0.055 | 6.31E-4 | -0.038 | 0.012 | -0.010 | 1.42E-5 | 3 | 1 | 1 | 0 |
| rs10401176 | 19 | 45253491 | T | C | BCL3 | intronic | -0.214 | 1.51E-11 | -0.160 | 2.46E-10 | -0.037 | 3.60E-30 | 14 | 1 | 1 | 0 |
| rs10401439 | 19 | 46320780 | T | C | SYMPK | intronic | 0.074 | 9.64E-6 | 0.070 | 5.50E-6 | 0.010 | 1.48E-5 | 8 | 1 | 4 | 1 |
| rs10406449 | 19 | 1040773 | A | G | ABCA7 | intronic | NA | NA | 0.082 | 0.038 | 0.018 | 0.001 | 70 | 1 | 6 | 1 |
| rs10409483 | 19 | 18590223 | G | C | ELL | intronic | 0.066 | 5.27E-5 | 0.062 | 3.38E-5 | 0.004 | 0.088 | 8 | 1 | 1 | 1 |
| rs10409727 | 19 | 45602862 | T | C | PPP1R37 | intronic | -0.072 | 0.001 | -0.080 | 4.03E-5 | -0.012 | 7.58E-5 | 43 | 1 | 1 | 2 |
| rs10409903 | 19 | 45600991 | C | T | PPP1R37 | intronic | -0.073 | 8.95E-4 | -0.081 | 3.70E-5 | -0.012 | 7.01E-5 | 40 | 1 | 1 | 2 |

Showing 1 to 10 of 304 entries    Previous 1 2 3 4 5 … 31 Next

## B.

Show 10 entries    Search: rs2280231

| SNP | CHR | POS | A1 | A2 | Gene | Location | IGAP_Beta | IGAP_Pvalue | Kunkle_Beta | Kunkle_Pvalue | Jansen_Beta | Jansen_Pvalue | Histone_mod | ATAC | diff_TF | eQTL |
|---|---|---|---|---|---|---|---|---|---|---|---|---|---|---|---|---|
| rs2280231 | 11 | 47600438 | T | C | KBTBD4 | UTR5 | 0.075 | 1.84E-5 | 0.067 | 1.99E-5 | 0.008 | 5.99E-4 | 58 | 32 | 40 | 3 |

Showing 1 to 1 of 1 entries (filtered from 304 total entries)    Previous 1 Next

**histone modifications**

| SNP | Tissue_Cell | Marker_name | ENCODE_ID |
|---|---|---|---|
| rs2280231 | angular gyrus | H3K27ac | ENCSR380KOO |
| rs2280231 | caudate nucleus | H3K27ac | ENCSR494MDB |
| rs2280231 | caudate nucleus | H3K27ac | ENCSR798RTU |
| rs2280231 | cingulate gyrus | H3K27ac | ENCSR604JDV |
| rs2280231 | cingulate gyrus | H3K27ac | ENCSR355UYP |
| rs2280231 | layer of hippocampus | H3K27ac | ENCSR912TVO |
| rs2280231 | layer of hippocampus | H3K27ac | ENCSR123HEE |
| rs2280231 | layer of hippocampus | H3K27ac | ENCSR321LKT |

**open chromatins (ATAC)**

| SNP | Tissue | Cell | Accession_ID |
|---|---|---|---|
| rs2280231 | Anterior cingulate cortex | glia | https://bendlj01.u.hpc.mssm.edu/multireg/ |
| rs2280231 | Anterior cingulate cortex | neuron | https://bendlj01.u.hpc.mssm.edu/multireg/ |
| rs2280231 | Amygdala | glia | https://bendlj01.u.hpc.mssm.edu/multireg/ |
| rs2280231 | Amygdala | neuron | https://bendlj01.u.hpc.mssm.edu/multireg/ |
| rs2280231 | Dorsolateral prefrontal cortex | glia | https://bendlj01.u.hpc.mssm.edu/multireg/ |
| rs2280231 | Dorsolateral prefrontal cortex | neuron | https://bendlj01.u.hpc.mssm.edu/multireg/ |
| rs2280231 | Hippocampus | glia | https://bendlj01.u.hpc.mssm.edu/multireg/ |
| rs2280231 | Hippocampus | neuron | https://bendlj01.u.hpc.mssm.edu/multireg/ |

**differential TF binding**

| SNP | Tissue/Cell | TF_name | ENCODE_Accession_ID | Pval_ref | Pval_snp | Pval_rank |
|---|---|---|---|---|---|---|
| rs2280231 | HepG2 | AGO2 | ENCFF622VSV | 0.010 | 0.020 | 0.049 |
| rs2280231 | GM12878 | ARID3A | ENCFF027VZK | 0.033 | 0.518 | 0.007 |
| rs2280231 | GM12878 | ARNT | ENCFF166OZW | 0.027 | 0.002 | 0.015 |
| rs2280231 | GM12878 | BHLHE40 | ENCFF095GMM | 0.036 | 0.002 | 0.002 |
| rs2280231 | K562 | BHLHE40 | ENCFF179NDS | 0.034 | 0.004 | 0.027 |
| rs2280231 | K562 | CREB3L1 | ENCFF582YPB | 0.004 | 0.150 | 0.020 |
| rs2280231 | HepG2 | CREM | ENCFF693JHE | 0.015 | 0.245 | 0.004 |

**eQTL**

| SNP | Dataset | Tissue_cell | Gene | beta | Pval |
|---|---|---|---|---|---|
| rs2280231 | eMeta | brain | MADD | 0.313 | 1.00E-8 |
| rs2280231 | eMeta | brain | MTCH2 | 0.407 | 0.00E+0 |
| rs2280231 | eMeta | brain | FNBP4 | 0.211 | 1.05E-5 |
| rs2280231 | 2017_NC_monocyte_ctl | monocyte | MTCH2 | 0.042 | 7.93E-4 |
| rs2280231 | 2017_NC_monocyte_ctl | monocyte | NUP160 | 0.055 | 5.41E-4 |
| rs2280231 | Raj_2014_monocyte | monocyte | NUP160 | -0.330 | 0.00E+0 |

**Supplementary Figure S2.** Annotation results of 304 functional variants at the AlzData (http://www.alzdata.org/functional_SNP_1.php). (A) Brief annotation results for 304 functional variants. (B) Detailed annotations for the query variant. CHR: chromasome; POS: position; A1: alternative allele; A2: reference allele; IGAP_beta: GWAS beta value from the Lambert study; IGAP_Pvalue: GWAS P value from the Lambert study [6]; Kunkle_beta: GWAS beta value from the Kunkle study; Kunkle_Pvalue: GWAS P value from the Kunkle study [7]; Jansen_beta: GWAS beta value from the Jansen study; Jansen_Pvalue: GWAS P value from the Jansen study [8]; Histone_mod: total number of datasets that had histone modification peaks (false discovery rate, FDR<0.001) overlapping with the target variant; ATAC: total number of assay for transposase-accessible chromatin with high throughput sequencing (ATAC-seq) datasets that had open chromatin peaks overlapping with the target variant (FDR<0.001); diffTF: total number of transcription factors (TFs) that could bind to the DNA element harboring the target variant (FDR<0.05) and binding affinities of these TFs with the DNA element were predicted to be significantly affected by different alleles of the target variant (P_rank<0.05); eQTL: expressional quantitive loci (eQTL) gene for the target variant ($P$<0.001).

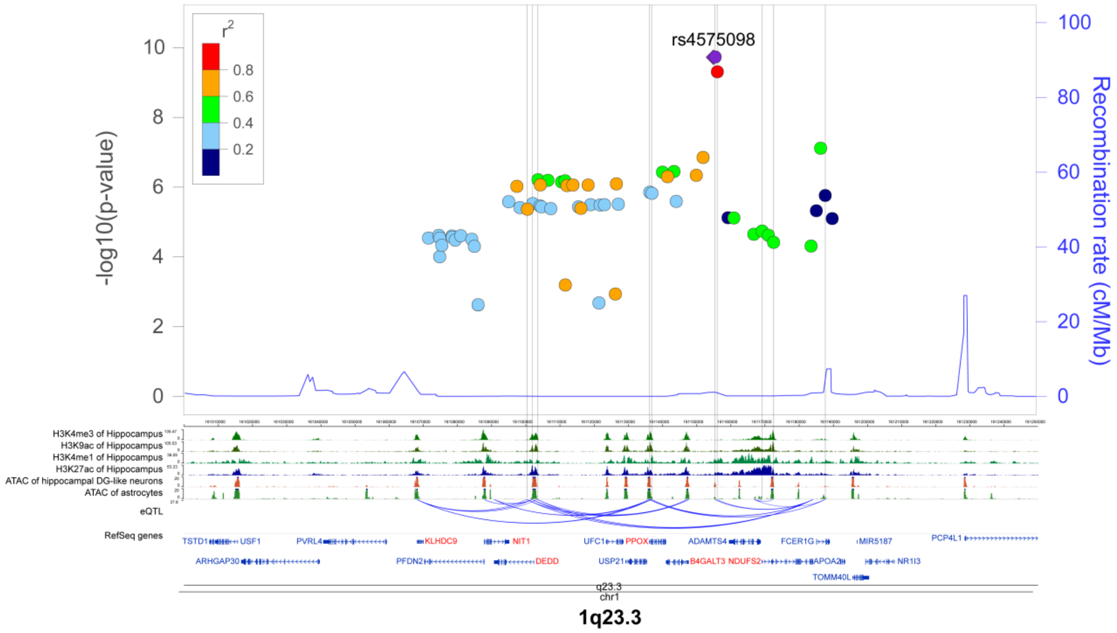


**Supplementary Figure S3**. Functional variants and causal genes identified in the 1q23.3 locus. (*Upper panel*) locus plot of GWAS results from the Jansen study [8]; (*lower panel*) functional annotation data including histone modification data from hippocampus, assay for transposase-accessible chromatin with high throughput sequencing (ATAC-seq) data from neurons and astrocytes, and eQTL data from brains and monocytes presenting the association among functional variants and causal genes; the locations of functional variants were shown in lightgrey lines and causal genes were marked in red.

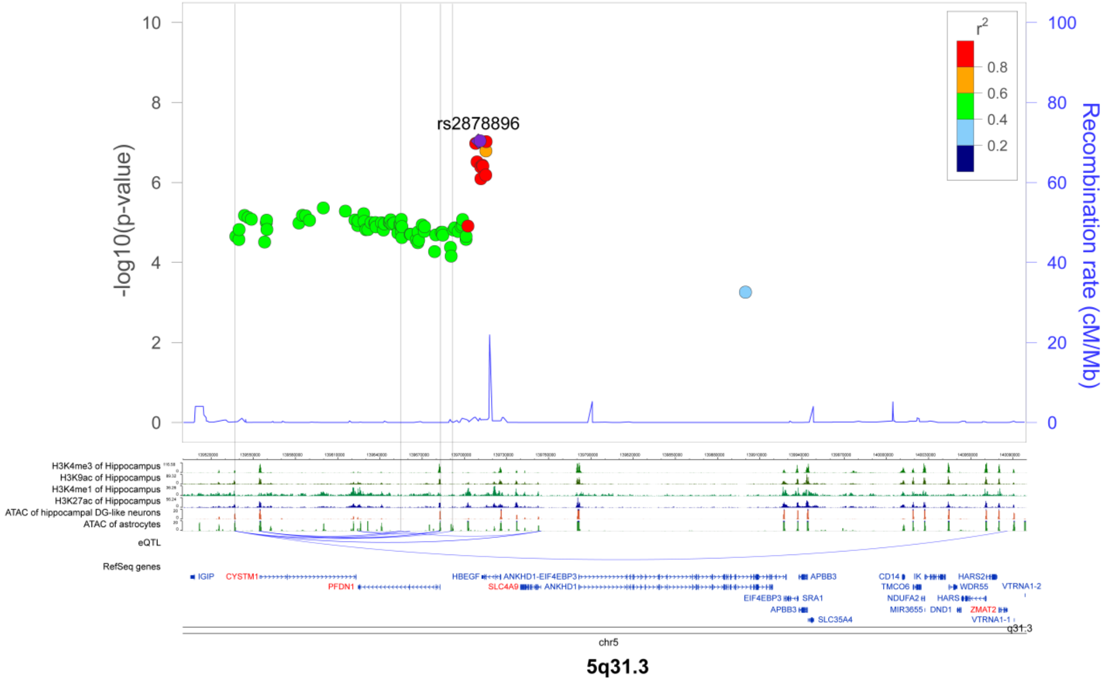


**Supplementary Figure S4**. Functional variants and causal genes identified in the 5q31.3 locus. (*Upper panel*) locus plot of GWAS results from the Lambert study [6]; (*lower panel*) functional annotation data including histone modification data from hippocampus, assay for transposase-accessible chromatin with high throughput sequencing (ATAC-seq) data from neurons and astrocytes, and eQTL data from brains and monocytes presenting the association among functional variants and causal genes; the locations of functional variants were shown in lightgrey lines and causal genes were marked in red.

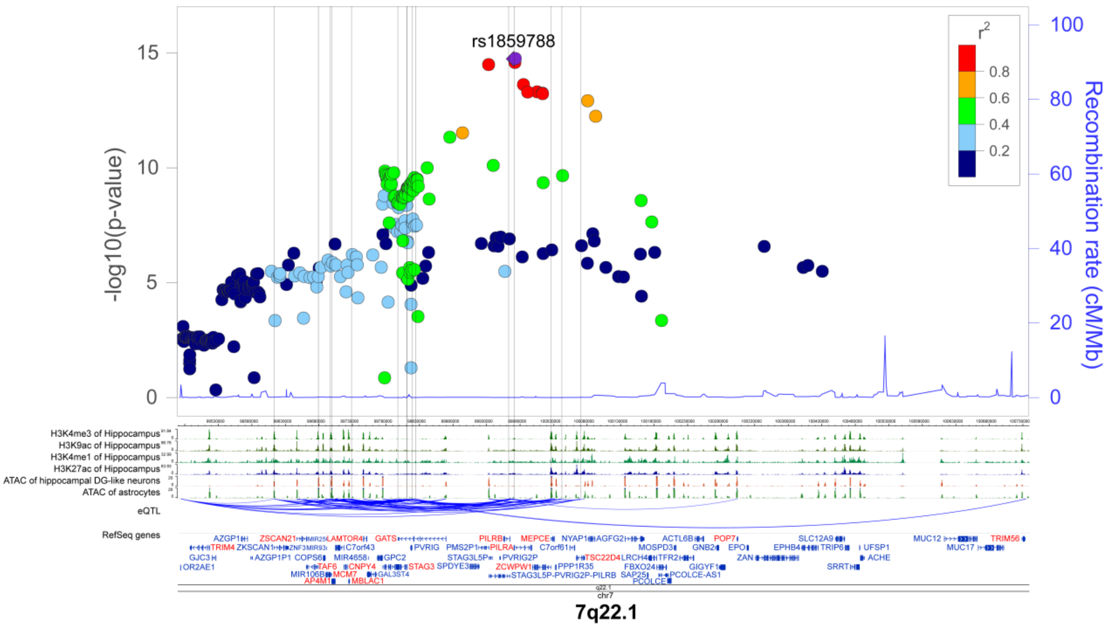


**Supplementary Figure S5**. Functional variants and causal genes identified in the 7q22.1 locus. (*Upper panel*) locus plot of GWAS results from the Jansen study [8]; (*lower panel*) functional annotation data including histone modification data from hippocampus, assay for transposase-accessible chromatin with high throughput sequencing (ATAC-seq) data from neurons and astrocytes, and eQTL data from brains and monocytes presenting the association among functional variants and causal genes; the locations of functional variants were shown in lightgrey lines and causal genes were marked in red.

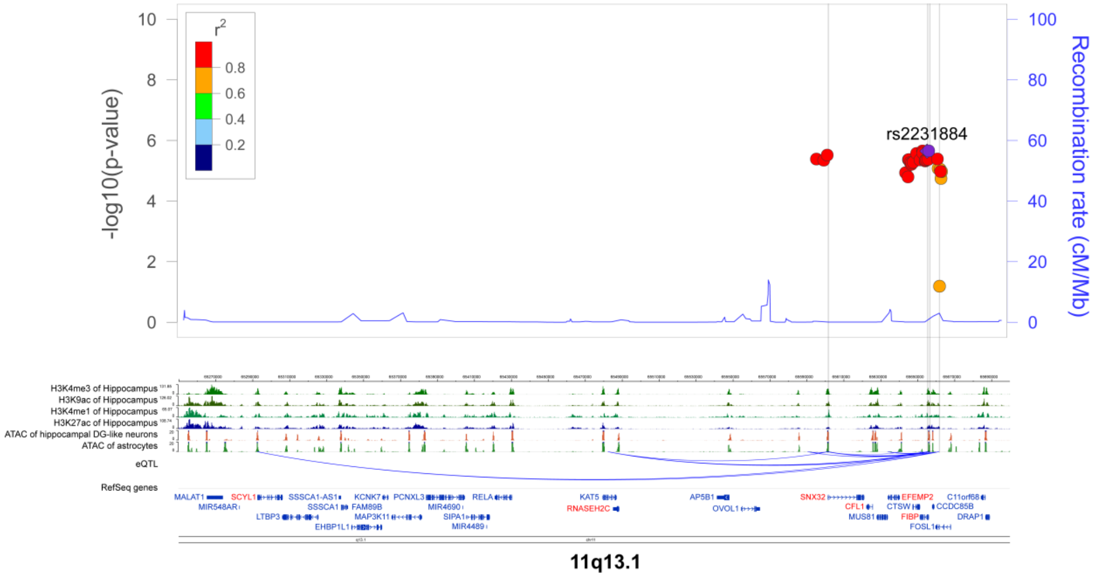


**Supplementary Figure S6**. Functional variants and causal genes identified in the 11q13.1 locus. (*upper panel*) locus plot of GWAS results from the Jansen study [8]; (*lower panel*) functional annotation data including histone modification data from hippocampus, assay for transposase-accessible chromatin with high throughput sequencing (ATAC-seq) data from neurons and astrocytes, and eQTL data from brains and monocytes presenting the association among functional variants and causal genes; the locations of functional variants were shown in lightgrey lines and causal genes were marked in red.

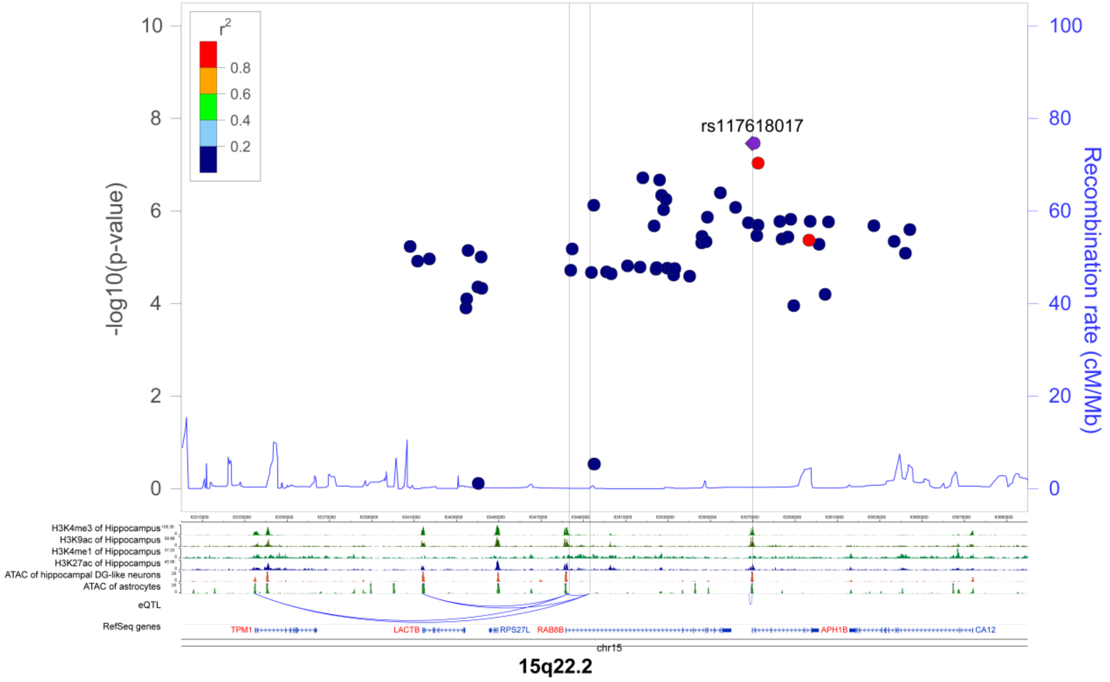


**Supplementary Figure S7**. Functional variants and causal genes identified in the 15q22.1 locus. (*upper panel*) locus plot of GWAS results from the Jansen study [8]; (*lower panel*) functional annotation data including histone modification data from hippocampus, assay for transposase-accessible chromatin with high throughput sequencing (ATAC-seq) data from neurons and astrocytes, and eQTL data from brains and monocytes presenting the association among functional variants and causal genes; the locations of functional variants were shown in lightgrey lines and causal genes were marked in red.

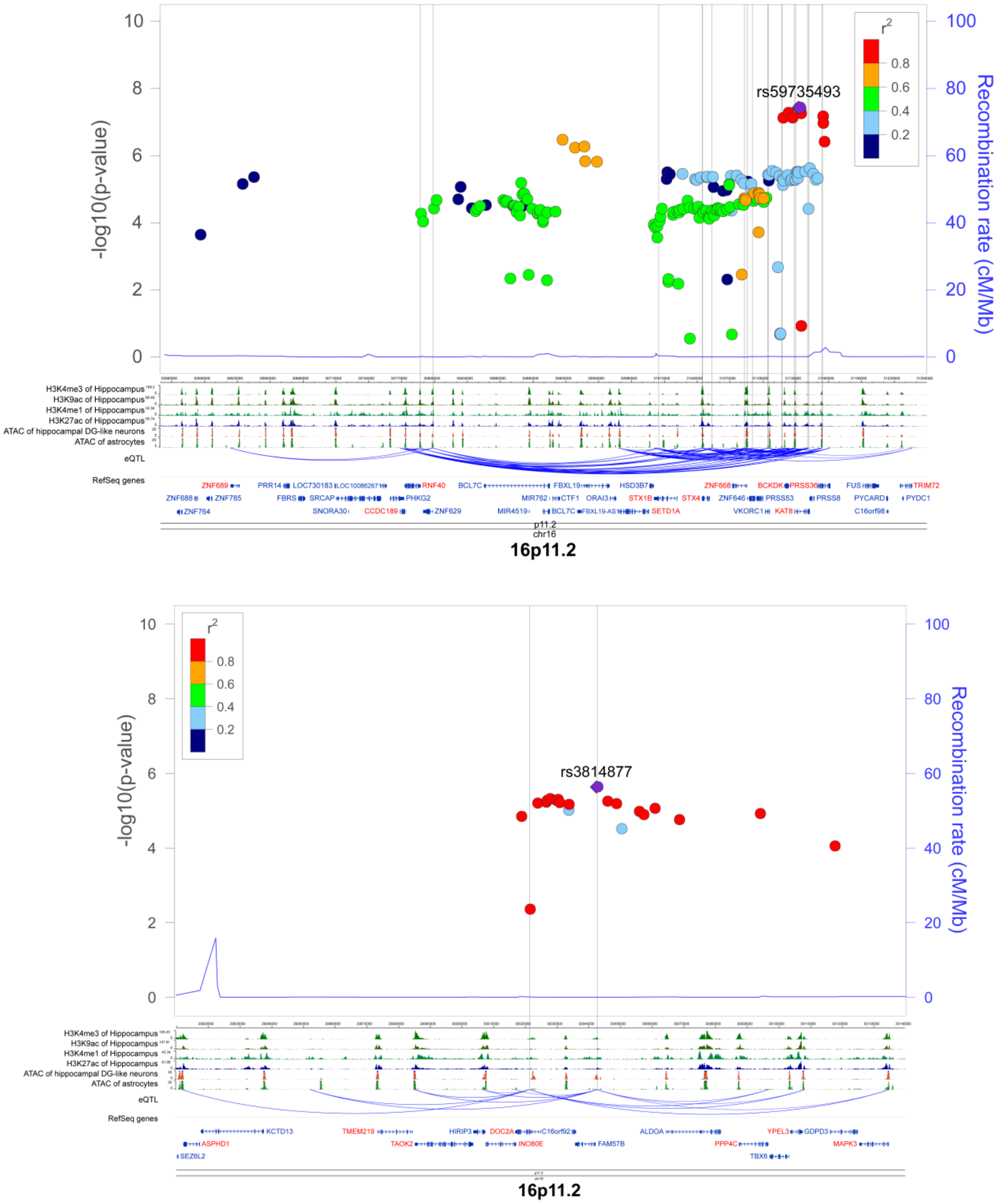


**Supplementary Figure S8**. Functional variants and causal genes identified in the 16p11.2 locus. (*upper panel*) locus plot of GWAS results from the Jansen study [8]; (*lower panel*) functional annotation data including histone modification data from hippocampus, assay for transposase-accessible chromatin with high throughput sequencing (ATAC-seq) data from neurons and astrocytes, and eQTL data from brains and monocytes presenting the association among functional variants and causal genes; the locations of functional variants were shown in lightgrey lines and causal genes were marked in red.

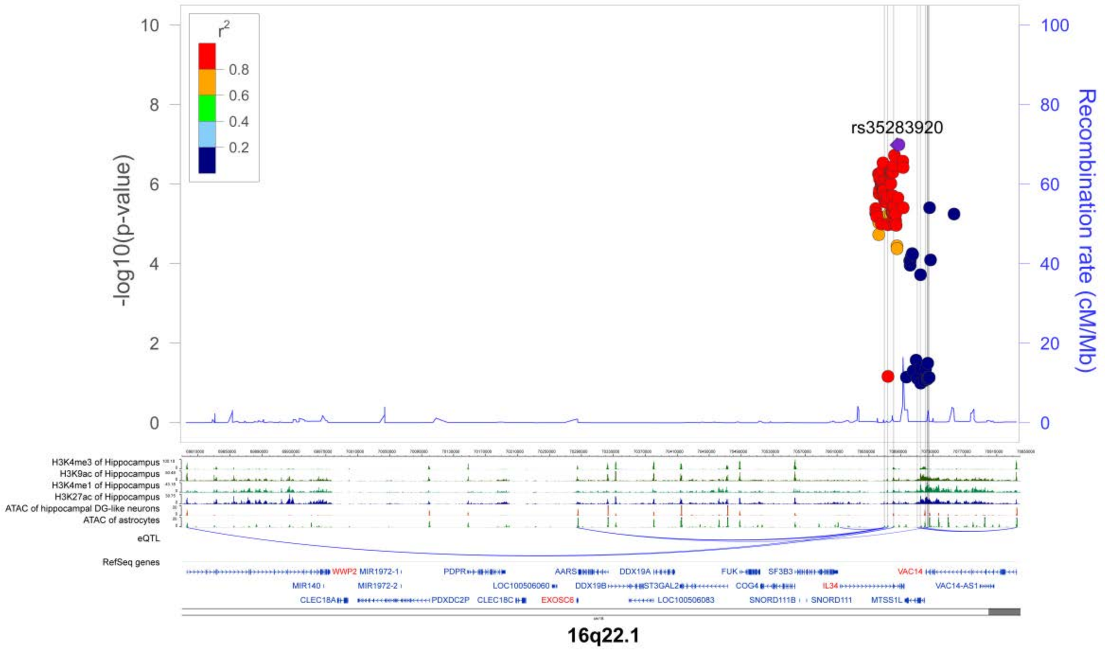


**Supplementary Figure S9**. Functional variants and causal genes identified in the 16q22.1 locus. (*upper panel*) locus plot of GWAS results from the Jansen study [8]; (*lower panel*) functional annotation data including histone modification data from hippocampus, assay for transposase-accessible chromatin with high throughput sequencing (ATAC-seq) data from neurons and astrocytes, and eQTL data from brains and monocytes presenting the association among functional variants and causal genes; the locations of functional variants were shown in lightgrey lines and causal genes were marked in red.

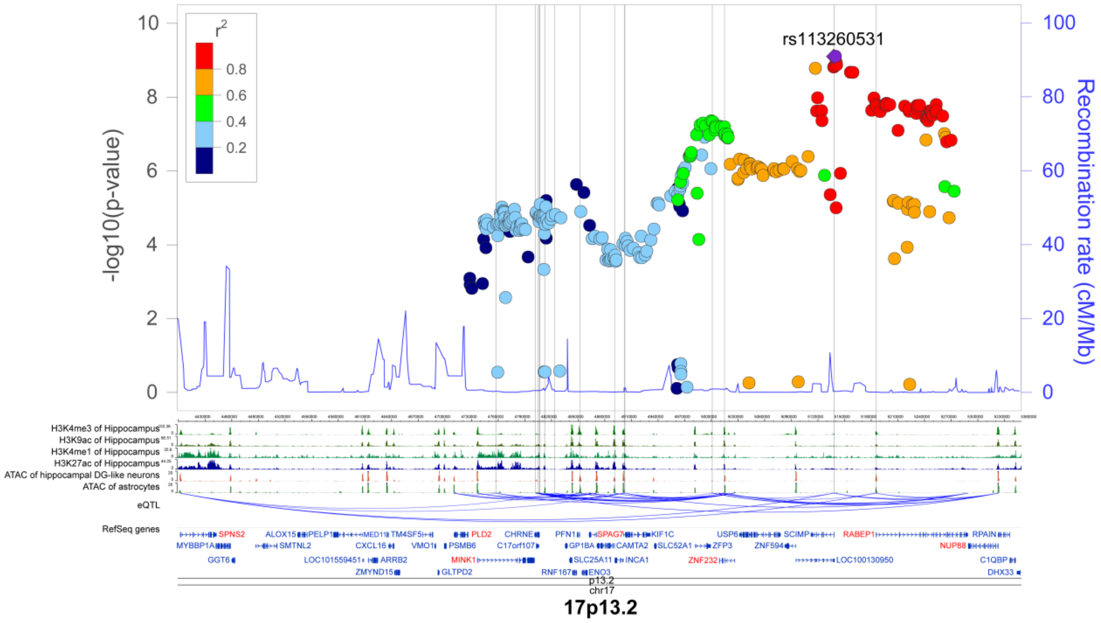


**Supplementary Figure S10**. Functional variants and causal genes identified in the 17p13.2 locus. (*upper panel*) locus plot of GWAS results from the Jansen study [8]; (*lower panel*) functional annotation data including histone modification data from hippocampus, assay for transposase-accessible chromatin with high throughput sequencing (ATAC-seq) data from neurons and astrocytes, and eQTL data from brains and monocytes presenting the association among functional variants and causal genes; the locations of functional variants were shown in lightgrey lines and causal genes were marked in red.

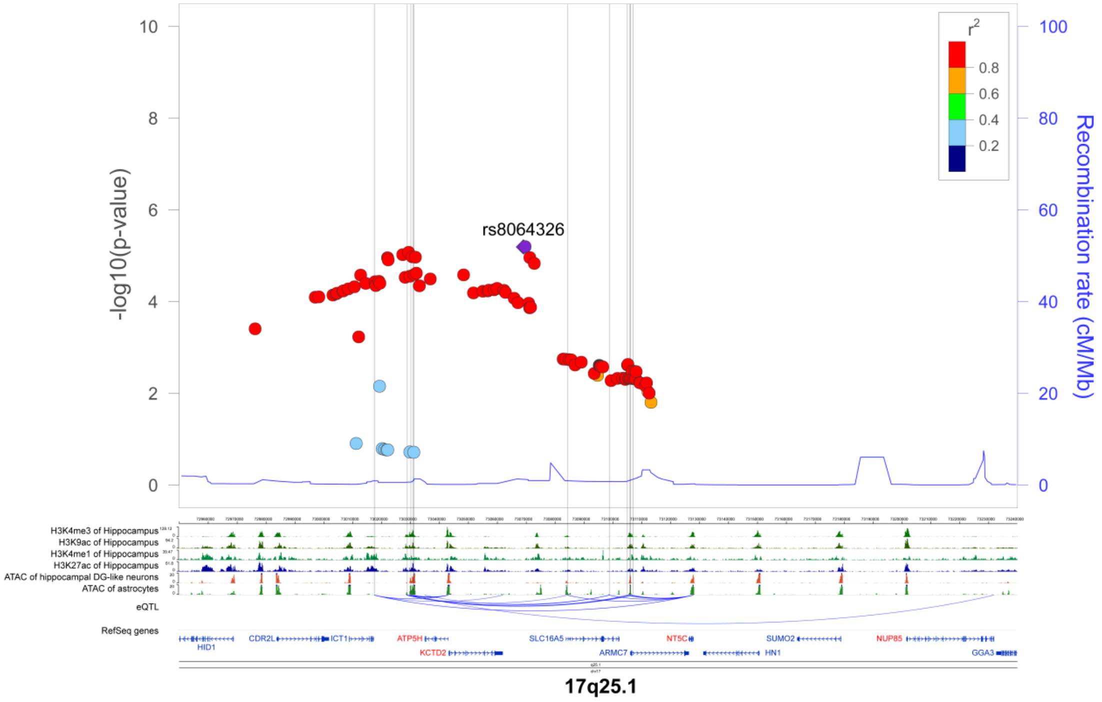


**Supplementary Figure S11**. Functional variants and causal genes identified in the 17q25.1 locus. (*upper panel*) locus plot of GWAS results from the Kunkle study [7]; (*lower panel*) functional annotation data including histone modification data from hippocampus, assay for transposase-accessible chromatin with high throughput sequencing (ATAC-seq) data from neurons and astrocytes, and eQTL data from brains and monocytes presenting the association among functional variants and causal genes; the locations of functional variants were shown in lightgrey lines and causal genes were marked in red.

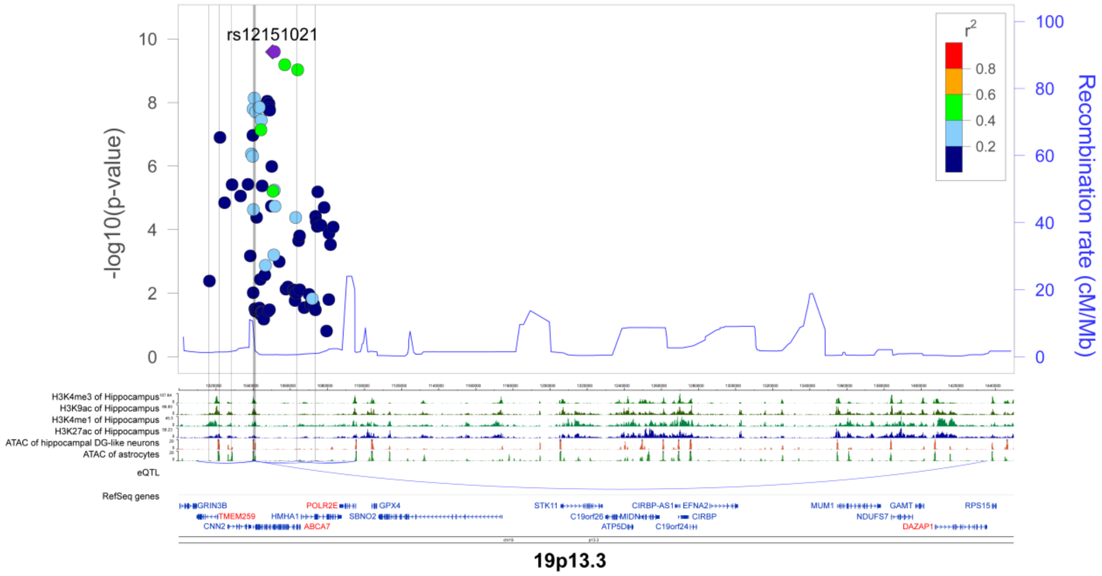


**Supplementary Figure S12**. Functional variants and causal genes identified in the 19p13.3 locus. (*upper panel*) locus plot of GWAS results from the Kunkle study [7]; (*lower panel*) functional annotation data including histone modification data from hippocampus, assay for transposase-accessible chromatin with high throughput sequencing (ATAC-seq) data from neurons and astrocytes, and eQTL data from brains and monocytes presenting the association among functional variants and causal genes; the locations of functional variants were shown in lightgrey lines and causal genes were marked in red.

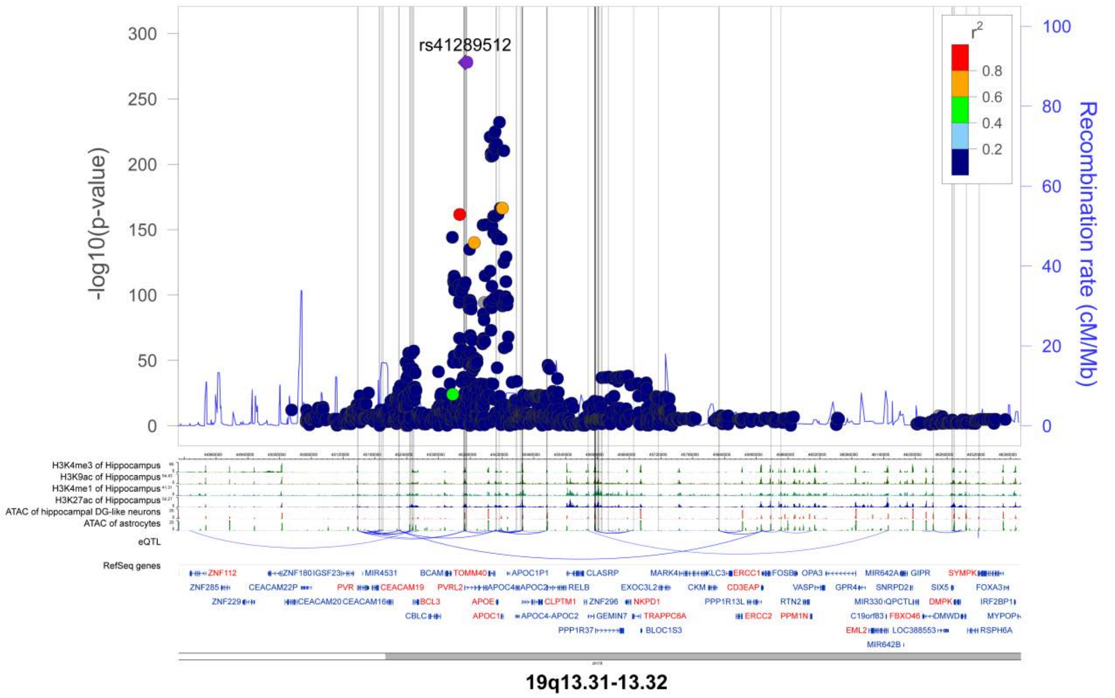


**Supplementary Figure S13**. Functional variants and causal genes identified in the 19q13.31-13.32 locus. **(***upper panel*) locus plot of GWAS results from the Jansen study [8]; (*lower panel*) functional annotation data including histone modification data from hippocampus, assay for transposase-accessible chromatin with high throughput sequencing (ATAC-seq) data from neurons and astrocytes, and eQTL data from brains and monocytes presenting the association among functional variants and causal genes; the locations of functional variants were shown in lightgrey lines and causal genes were marked in red.

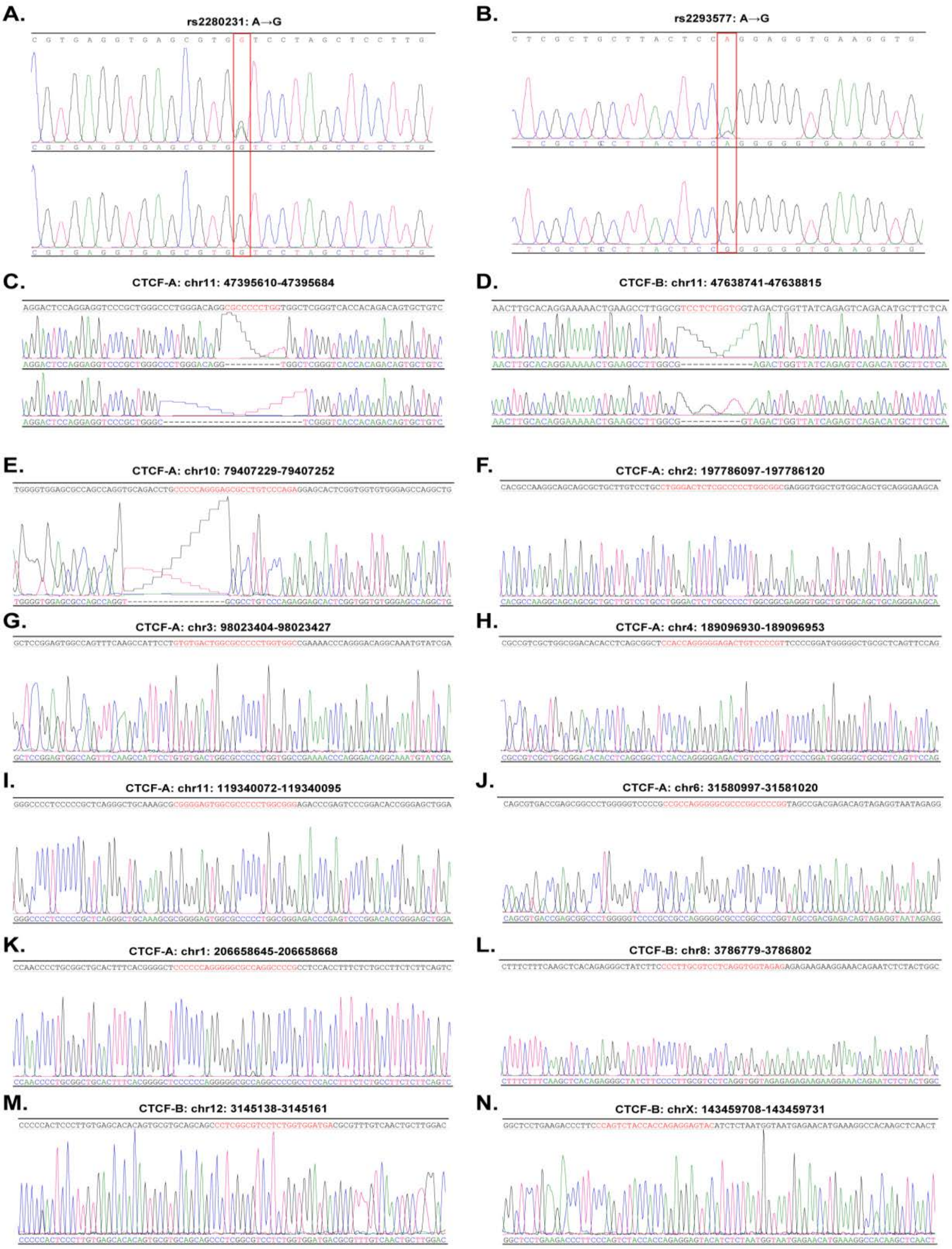


**Supplementary Figure S14.** Sanger sequencing results. Successful base-editing of (A) rs2280231 and (B) rs2293577 in 293T cells. Highlighted bases were targeted SNPs for A to G edition. Successful knockout of CTCF-binding sites in U251-APP cells for (C) CTCF-A and (D) CTCF-B. Sequencing of predicted off-target sites in (E~K) CTCF-A and (L~N) CTCF-B cells. Regions highlighted with red were predicted off-target sites.

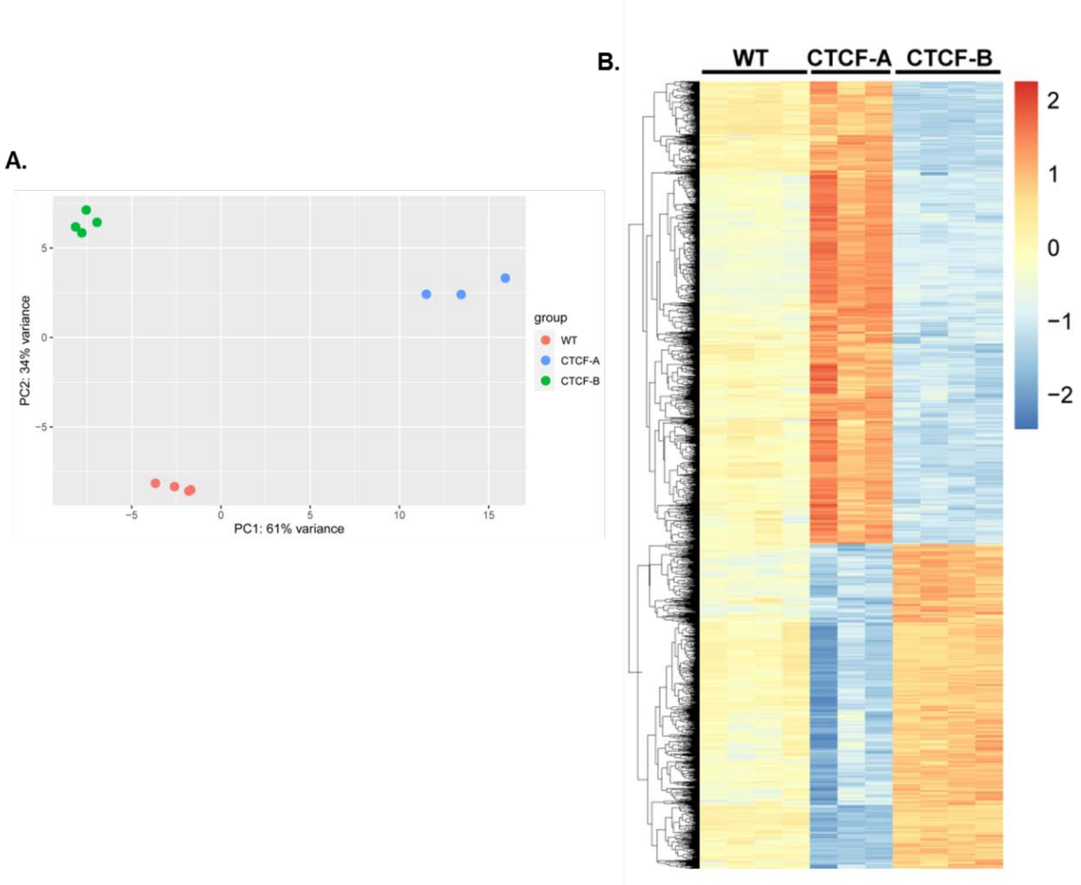


**Supplementary Figure S15.** RNA sequencing results for wild-type U251-APP cell (WT), CTCF-A cell, and CTCF-B cell. (A) PCA plot and (B) Heatmap for common differentially expressed genes in CTCF-A cell and CTCF-B cell compared to WT.
